# Fibrillarin shapes oncogenic protein pools and ribosomal composition in triple-negative breast cancer

**DOI:** 10.1101/2025.03.21.644506

**Authors:** Paula Groza, Kanchan Kumari, Johanna Schott, Margalida Esteva-Socias, Eliana Destefanis, Carlos Peula, Chloe Williams, Virginie Marchand, Pernilla Wikström, Rebecca Wiberg, Ana Bosch Campos, Jonathan D. Gilthorpe, Bogdan Pop, Andre Mateus, Yuri Motorin, Erik Dassi, Francesca Tuorto, Francesca Aguilo

## Abstract

Fibrillarin (FBL), a core component of the C/D box snoRNP complex, catalyzes 2’-O-methylation (Nm) of ribosomal RNA (rRNA), influencing ribosome heterogeneity and oncogene translation. In triple-negative breast cancer (TNBC), FBL dysregulation creates an aberrant Nm signature. This study explores role of FBL in TNBC progression via translation-driven mechanisms. FBL knockdown impaired tumorigenic traits, induced metabolic stress, and reduced the translation efficiency of oncogenes, including metastasis-associated protein 1 (MTA1), interleukin-1 receptor-associated kinase 1 (IRAK1), and thymosin beta 10 (TMSB10). RiboMethSeq analysis revealed differential rRNA Nm site sensitivity to FBL depletion. Additionally, FBL knockdown lowered RPS28 protein levels, suggesting its misincorporation into ribosomes. Notably, silencing RPS28 also suppressed oncogenic traits and downregulated MTA1, IRAK1, and TMSB10, highlighting its role in FBL-mediated translation. These findings uncover a complex interplay between FBL, rRNA Nm modifications, and RPS28 in shaping oncogenic protein pools and ribosomal composition in TNBC. Targeting this pathway could offer novel therapeutic strategies for this aggressive cancer subtype.

## Introduction

In cancer, gene expression is altered at multiple levels, and deregulated translation plays a key role in sustaining tumorigenesis by driving enhanced proliferation and growth rates. Regulation of translation involves chemical modifications of various RNA species, including ribosomal RNA (rRNA) and messenger RNA (mRNA) ^1,2^. RNA modifications are key regulators of gene expression and numerous studies have linked them to the development of various pathologies, including cancer ^3–5^. Among these, 2’-O-methylation (Nm) is the most abundant rRNA modification and involves the addition of a methyl group to the 2’ hydroxyl of the ribose moiety in a nucleoside ^6^. Nm is essential for rRNA folding, stability, catalytic activity, and interactions with antibiotics and other molecules such as tRNAs, mRNAs, and proteins ^6^.

This modification is catalyzed by a small nucleolar ribonucleoprotein (snoRNP) complex comprising a site-specific C/D box small nucleolar RNA (SNORD) and four ribonucleoproteins: NOP56, NOP58, SNU13 (or NHP2L1), and fibrillarin (FBL) ^7,8^. snoRNAs ensure substrate specificity through base pairing with rRNA, and FBL catalyzes the deposition of Nm at specific sites ^9^. Aberrant Nm levels and altered expression of FBL or SNORDs have been linked to changes in translation efficiency and fidelity, contributing to tumor progression in multiple cancer types, including breast cancer ^10–12^. Triple-negative breast cancer (TNBC), a heterogeneous and aggressive subtype of breast cancer, is characterized by a lack of the estrogen receptor (ER), progesterone receptor (PR), and human epidermal growth factor receptor 2 (HER2), making it unsuitable for hormonal therapies ^13^. Genomic and transcriptomic studies have categorized TNBC into four distinct molecular subtypes with unique clinical outcomes ^14,15^, highlighting the need for tailored treatments. Therapeutic approaches targeting oncogenic translation processes involved in metastasis, drug resistance, and tumor progression hold significant promise for improving patients’ TNBC outcomes.

Recent findings suggest that heterogeneous ribosomes play a critical role in driving oncogenic translation and enhancing tumorigenic potential ^16–18^. Ribosome heterogeneity arises from ribosomal DNA (rDNA) variants, altered RNA Polymerase I activity, changes in ribosomal protein expression, and alterations in rRNA modifications ^19^. Studies have shown that oncogenic ribosomes present differential Nm levels at specific rRNA sites compared with normal cells ^20,21^. For instance, increased Nm at position A576 and decreased Nm at position G1447 on 18S rRNA have been identified in TNBC cell lines and human samples, although the functional impact of these changes remains unclear ^22^.

In the present study, we investigated the role of FBL in driving oncogenic translation in TNBC cells. FBL silencing significantly reduced the proliferation and migration of MDA-MB-231 TNBC cells, altered their translational landscape, and induced metabolic stress. Notably, FBL depletion decreased the translation of TNBC-associated oncoproteins such as metastasis-associated protein 1 (*MTA1*), interleukin-1 receptor-associated kinase 1 (*IRAK1*), and thymosin beta 10 (*TMSB10*), a phenomenon specific to TNBC cells that was not observed in healthy epithelial cells. Using RiboMethSeq, we showed that Nm sensitivity to FBL varied across sites. Furthermore, FBL silencing influenced ribosome composition, specifically affecting the incorporation of the ribosomal protein RPS28. The loss of RPS28 also impaired the tumorigenic properties of TNBC cells and decreased the translation of MTA1, IRAK1, and TMSB10, underscoring its critical role in oncogenic translation. Collectively, these findings identify FBL, rRNA Nm, and RPS28 as critical drivers of oncogenic translation in TNBC, and offer potential therapeutic targets for this aggressive cancer.

## Materials and methods

### Antibodies

The following antibodies were used at the indicated concentrations: anti-FBL (A303-891A, Bethyl Laboratories, 1:2,500 dilution), anti-β-actin (A5441, Sigma, 1:10,000 dilution), anti-metastasis-associated 1 gene (MTA1) (A300-911A-T, Bethyl Laboratories, 1:2,000), anti-interleukin 1 receptor-associated kinase 1 (IRAK1) (ab302554, Abcam, 1:2,000 dilution), anti-thymosin B10 (sc-514309, Santa Cruz Biotechnology, 1:1,000 dilution), anti-RPS28 (14796-1-AP, Proteintech, 1:2,000 dilution), and anti-RPL22 (25002-1-AP, Proteintech, 1:2,000). The secondary antibodies used were Goat Anti-Rabbit IgG H&L horseradish peroxidase (HRP) (Abcam, ab6721, 1:10,000) and Goat Anti-Mouse IgG H&L (HRP) (Abcam, 1:10,000, ab6789).

### Cell culture and antibiotic selection

MDA-MB-231 cells were maintained in Dulbecco’s Modified Eagle’s medium (DMEM) enriched with 10% fetal bovine serum (FBS), 1% penicillin/streptomycin, and 0.1% human insulin. hTERT-HME1 cells were cultured in F12/DMEM supplemented with 5% horse serum, 0.1% insulin, cholera toxin, human epidermal growth factor, hydrocortisone, and 1% penicillin-streptomycin.HEK-293T cells were cultured in DMEM supplemented with 10% FBS and 1% penicillin-streptomycin. All cells were maintained in a humidified incubator at 5% CO_2_ and 37 °C. Puromycin (1 µg/ml) and hygromycin (500 µg/ml) were used to select the transduced cells.

### Protein extraction and Western Blot analysis

Cells were harvested and lysed using whole-cell extract lysis buffer (50 mM HEPES pH 7.5, 150 mM NaCl, 3 mM MgCl2, 0.2% Triton X-100, 0.2% Nonidet NP-40, 10% glycerol, and 1X protease inhibitor cocktail). For Western Blot analysis, equal amounts of protein extracts were subjected to SDS-PAGE and transferred to nitrocellulose or PVDF membranes using wet transfer. The membranes were blocked by incubation in 5% skim milk dissolved in PBS-T (1X PBS, 0.1% Tween-20) for 1 h at room temperature (RT). Primary antibodies were incubated overnight at 4⁰C with slight shaking at the dilutions above. After washing, the membranes were incubated with secondary antibody for 1 h at RT. The signal was detected using Pierce^TM^ ECL Western Blotting substrate (Thermo Scientific™) using Amersham™ Imager 680.

### Generation of knockdown by short hairpin RNA

To generate gene knockdowns, we used two distinct short hairpin RNAs (shRNAs) for each of the targeted genes (*FBL* and *RPS28*) (**Table S1**), which are later referred to as KD1 and KD2. The shRNAs were cloned into pLKO.1-Puro and pLKO.1-Hygro (Addgene #8453 and #24150) following Addgene recommendations, with a few modifications. Briefly, the plasmids were digested using AgeI and EcoRI restriction enzymes (Anza, Invitrogen). shRNA oligos were annealed and ligated into the pLKO.1 backbone vector. The cloning was validated by Sanger sequencing. Lentiviral particles were generated by transfecting HEK-293T cells with lentiviral plasmids pLKO.1-Puro/Hygro containing shRNA or scramble (Scr) control (8 µg), pCMV-dR8.2 packaging (6 µg), or pCMV-VSV-G helper (2 µg) plasmids using the CaCl_2_ method. Briefly, the plasmid DNA was mixed with 2M CaCl_2_ and MQ water. The mixture was added to 2X HEPES Buffer Solutions (HBS), vortexed, incubated for 20 min at RT, and added to the media. Lentiviral particles were collected after 48 and 72 h of incubation and concentrated using 30 kDa Amicon Ultra-15 Centrifugal Filters (Merck). MDA-MB-231 cells were transduced with lentiviral particles in a medium supplemented with polybrene (8 µg/ml). After incubation with the virus for 48 h, the cells were selected with either 1 µg/ml puromycin or 500 µg/ml hygromycin for 48 or 96 h, as indicated.

### RNA extraction, first-strand cDNA synthesis, and Real-time quantitative PCR (RT-qPCR)

Total RNA was isolated and extracted using the RNeasy® Plus Mini Kit (Qiagen®) or Qiazol, following the manufacturer’s protocols, and eluted in 30 µL of nuclease-free water. RNA was extracted from the polysome fractions using the phenol:chloroform:isoamyl alcohol (25:24:1, pH 4,5-5) method using ROTI®Aqua-P/C/I (CarlRoth®). First-strand cDNA synthesis was performed using a RevertAid® First Strand cDNA Synthesis Kit (Thermo Scientific ™). Reverse transcription (RT) followed by PCR was performed according to the manufacturer’s protocol, using random primers from the kit. Real-time quantitative PCR (RT-qPCR) was performed using SYBR™ Green PCR Master Mix (Applied Biosystems™), according to the manufacturer’s protocol. The primers used for RT-qPCR are listed in **Table S1**.

### MTT assay

After 96 h of selection with puromycin, control and KD MDA-MB-231 cells were seeded in 96-well plates (3 × 10^3^ cells/well) and incubated at 37 °C. MTT (3-(4,5-dimethylthiazol-2-yl)-2,5-diphenyltetrazolium bromide) (Sigma-Aldrich) was added at a final concentration of 0.5 mg/ml at different time points after seeding and incubated for 4 h at 37 °C. The medium containing MTT was aspirated, formazan crystals were dissolved in DMSO, and the absorbance at 570 nm was measured using a Tecan Spark® multimode microplate reader.

### Calcein staining

An equal number of control and FBL KD MDA-MB-231 cells were seeded in a 96-well plate (2 × 10^4^ cells/well) following selection with puromycin. The following day, calcein AM (Invitrogen) was added to each well at a final concentration of 5 µg/ml, and the cells were incubated for 30 min at 37 °C. Thereafter, 10 µL of Hoechst 33342 (5 μg/ml, Thermo Fisher Scientific^TM^) was added to each well and the plate was incubated for an additional 10 min.

The medium was then replaced with 200 µL of serum-free media containing 15 µL of hemoglobin, and images were taken using a Tecan Spark Cyto® multimode microplate reader with excitation/emission settings of 405 nm/450 nm for Hoechst and 495 nm/515 nm for calcein, at 10X magnification. Quantification of calcein-positive cells was performed using the Fiji (ImageJ) software. For each experimental condition, three non-overlapping areas (869 × 640 pixels) were analyzed for three distinct images. The stained cells in the area were counted using the counting tool from ImageJ. The results were averaged across three biological replicates.

### Crystal violet assay

Control and KD MDA-MB-231 cells were seeded in duplicates in 6 well-plates at a density of 5 × 10^3^ cells/well following puromycin selection and maintained in culture for 10 days, with growth media refreshed every 2 days. At the end of the incubation, the medium was removed, and the cells were washed with 1X PBS. Subsequently, they were fixed with 10% formaldehyde, stained with crystal violet for 1 h at RT, and washed three times with 1X PBS.

### Wound healing

The migration capacity of control and KD MDA-MB-231 cells was assessed using a wound healing assay. Following the 96 h period of selection with puromycin, cells were seeded into 12-well plates and incubated at 37 °C for 24 h to achieve full confluency. A wound was scratched using a sterile 200 µl pipette tip. Images were captured using a Nikon Eclipse Ts2 microscope at specific time points, that is, immediately after the scratch and 24 h later. Wound closure was quantified using Fiji (ImageJ) software.

### Transwell assay

The transwell migration assay for control, FBL, and RPS28 KD MDA-MB-231 cells was performed using cell culture inserts with 8 µm pores (Falcon, Corning®) in a 24-well plate format. Briefly, 1 × 10^5^ cells were seeded into the transwell chamber in 100 µL DMEM without FBS, and 600 µL of complete medium was added to each well containing a transwell. After 24 h, the transwells were removed and fixed with 10% formaldehyde for 20 min. After fixation, the transwells were washed with 1X PBS and dried for 10 min. Non-migrating cells were gently removed from the upper surface of the transwell membrane using a cotton-tipped applicator. To stain the migrated cells, 600 µL of 0.2% crystal violet was added to each well of the 24-well plate, and each transwell was immersed in the staining solution and incubated for 30 min at RT. The transwells were washed with 1X PBS 3 times and dried overnight at RT. Images were captured using an EVOS XL Core Imaging System at 10X magnification. Fiji (Image J) software was used to quantify the area occupied by the migrated cells, which was then used to calculate the relative migration capacity.

### Apoptosis

To assess apoptosis, 1 × 10^5^ control and FBL KD MDA-MB-231 cells were collected, resuspended in 1X binding buffer, and stained with 7-Aminoactinomycin D (eBioscience^TM^ 7-AAD Viability Staining, Invitrogen) and Annexin V (Alexa Fluor^TM^ 488 Ready Flow^TM^, Invitrogen). Following a 15 min incubation at RT, the cells were immediately analyzed using an Accuri C6+ Flow cytometer (BD Biosciences).

### Polysome profiling

Control and FBL/RPS28 KD MDA-MB-231 cells were treated with cycloheximide (CHX) (100 µg/ml) for 5 min and washed twice with cold 1X PBS containing CHX (100 µg/ml). The cells were then trypsinized, resuspended in complete medium with CHX (100 µg/ml), and counted. Equal numbers of cells were lysed in polysome lysis buffer (20 mM Tris-HCl, pH 7.4, 5 mM MgCl_2_, 150 mM NaCl, 1% Triton X-100) supplemented with CHX (100 µg/ml), 1X complete protease inhibitors (complete EDTA-free, Roche), and 200 U/ml of RiboLock RNase Inhibitor (Thermo Scientific). Lysates were tumbled for 10 min and then centrifuged at 10,000 *g* for 10 min at 4 °C. Cleared lysates were layered on sucrose gradients (15-45% w/v) created using a Gradient Master (BioComp Instruments). The samples were centrifuged at 39,000 *g* for 2 h at 4 °C in an ultracentrifuge SW60 Ti rotor head (Beckmann). The absorbance at 260 nm (A260) was continuously recorded using a Triax™ flow cell to generate polysome profiles (BioComp Instruments). Fractions of 280 µL were collected using a Gilson Fraction FC-203B collector in tubes containing 300 µL urea solution (10 mM Tris-HCl pH 7.5, 350 mM NaCl, 10 mM EDTA, 1% SDS, and 42% urea). Absorbance values were plotted against the piston position (1.00 - 54.00 mm) to generate polysome profiles.

For each experimental condition, 14 fractions were collected and total RNA was extracted, followed by first-strand cDNA synthesis and qPCR. The distribution of mRNA among the fractions from the FBL KD experiment was calculated using cycle threshold (Ct) values. ΔCt for each fraction was calculated as Ct(fraction 1)–Ct (fraction). ΔCt was then used to calculate 2^-ΔCt^, which was further summed for all fractions, and the percentage of mRNA was calculated as (2^-ΔCt(fraction^ ^n)^ × 100)/ sum of all 2^-ΔCt^. For RPS28 KD, the fractions were combined for qPCR as subpolysomal fractions (fractions 1–6) and polysomal fractions (fractions 7–14). The mRNA distribution of RPS28 KD was calculated relative to the subpolysomal fractions of Scr.

### Ribosome profiling (Ribo-seq)

For Ribo-seq, cytoplasmic lysates from control and FBL-KD MDA-MB-231 cells were collected as described in the polysome profiling section. For each sample, the absorbance of the lysate was measured using a NanoDrop spectrophotometer, and 10 µg of RNA was stored as the input cytoplasmic RNA. The remaining lysate was treated with 4 U DNase I (Thermo Scientific) and 800 U RNase I (Ambion) for 45 min at RT with gentle shaking. 800 U of RNasin ribonuclease inhibitor (Promega) was added to quench the reaction, and the samples were run on a 17.5 – 50% sucrose gradient to isolate monosomes. After separation through ultracentrifugation, the gradients were fractionated using an ISCO gradient fractionator (Brandel), and the fractions were collected in 300 µL of urea solution (see the polysome profiling section). RNA from the monosome-containing fraction and input samples was isolated using phenol:chloroform:isoamyl alcohol (25:24:1). RNA was quantified using the 2200 Agilent R6K ScreenTape station. Monosome-protected RNA fragments were dephosphorylated using T4 polynucleotide kinase (PNK) (Takara), and RNA was extracted using acid-phenol. RNA was separated on a 15% polyacrylamide TBE-urea gel, and footprint fragments of 20-30 nucleotides in length were isolated from the gel. Sequencing libraries were prepared according to the protocol of the NEB NEXT Small RNA Library Prep Set for Illumina (Multiplex Compatible) E7330 Kit. The cytoplasmic RNA (RNA Input) was sequenced using Illumina HiSeq V4 SR 50, whereas the footprint library used small RNA Seq-HiSeq V4 SR 50. A fold-change threshold of ±1.5 (log2 transformed) was applied to identify the downregulated and upregulated genes.

### Processing of Ribo-seq reads

The samples were sequenced using the NEXTSeq550 system. After sequencing, the adapters of the ribosome footprint sequences (AGATCGGAAGAGCACACGTCT) were removed using the FASTX-Toolkit (http://hannonlab.cshl.edu/fastx_toolkit/), retaining only sequences of at least 20 nucleotides in length. Alignment was performed using Bowtie v1.2.2 ^23^, allowing a maximum of two mismatches and reporting all alignments in the best stratum (settings: -a –best –stratum -v 2). Reads that were mapped to human tRNA (downloaded from GtRNAdb) or rRNA sequences (downloaded from the UCSC Genome Browser) were removed. The remaining reads were aligned to the human transcriptomic sequences (obtained from the UCSC table wgEncodeGencodeBasicV38). Next, reads were summarized at the gene level using Samtools v1.2 ^24^. An in-house developed Perl Script was used to count only reads between 26 and 30 nucleotides, which were mapped to ORFs of isoforms arising from one specific gene (as defined by a common gene symbol). An offset of 12 nucleotides upstream of the start codon and 15 nucleotides upstream of the stop codon for the 5’ end of the reads was assumed. RNA-seq reads of the input samples (80 nucleotide single-end) were processed analogously, omitting adapter removal and size selection. Finally, 1.7 to 6.1 million reads could be uniquely assigned to genes in each footprint sample, and 23.5 to 27.6 million reads in each input sample. After generating the aligned reads, codon usage analysis was performed using the RiboView tool as previously described ^25^. For ribosome differential composition, the differential ribosomal protein incorporation prediction by analysis of rRNA fragment (dripARF) software tool was used ^26^.

### Ribo-seq output and general quality control

The periodicity of the reads was assessed by plotting the distribution of the 5’ end of all uniquely assigned reads relative to the start and stop codons. The strongest periodicity was observed for 28 nucleotide-long footprint fragments. After adapter removal, 28–29 nucleotide-long reads were the most abundant fragment size among the footprint reads. Ribosome densities were calculated from read counts using DESeq2 v1.34.0 ^27^ for normalization (applying the median of ratios method) and checking for statistically significant differences after KD compared to control cells (with a Likelihood Ratio Test followed by the Benjamini-Hochberg procedure for multiple testing adjustment). The translation efficiency of specific mRNAs was calculated as the ratio of their Ribo-seq reads (actively translated mRNAs) to RNA-seq reads (total mRNA abundance). Transcripts were further separated into different categories (forwarded, buffered, intensified, and exclusive) based on the changes in RNA abundance, ribosome protected fragments (RPFs), and translation efficiency using deltaTE software as previously described ^28^.

### Mass spectrometry-based proteomics

To assess protein expression, mass spectrometry-based proteomics was employed. 10 μg of protein from control and FBL KD MDA-MB-231 cells were reduced to a final concentration of 20 mM tris(2-carboxyethyl) phosphine (TCEP) before being digested with a modified sp3 protocol, as previously described ^29^. Briefly, a bead suspension (10 μg of beads (Sera-Mag Speed Beads, 4515-2105-050250, 6515-2105-050250) in 10 μl 15% formic acid and 30 μl ethanol) was added to the samples, followed by incubation for 15 min at RT with shaking, and the beads were washed four times with 70% ethanol. Proteins were digested overnight by adding 40 μl of digestion solution (5 mM chloroacetamide, 1.25 mM TCEP, and 200 ng trypsin in 100 mM HEPES pH 8.5). After elution from the beads, the peptides were dried under vacuum and labeled with TMT10plex (Thermo Fisher Scientific). After pooling, the samples were desalted by solid-phase extraction using a Waters OASIS HLB μElution plate (30 μm). Finally, the samples were fractionated into 48 fractions on a reversed-phase C18 system under high pH conditions, pooling every twelfth fraction. Samples were analyzed by LC-MS/MS using a data-dependent acquisition strategy on a Thermo Fisher Scientific™ Vanquish Neo LC coupled with a Thermo Fisher Scientific^TM^ Orbitrap Exploris 480. Raw files were processed with MSFragger ^30^ against the human FASTA database downloaded from UniProt (UP000005640) using standard settings for TMT. To normalize the data across conditions, we first identified 200 proteins with the highest fold change (FC) between each replicate of the FBL KD and control conditions. Since translation was impaired, we assumed these proteins did not change their levels. Thus, we fitted a vsn model to these proteins ^31^ and applied it to the entire dataset. We calculated the median intensity for each condition and the overall median, and determined the difference. Finally, a correction factor was applied to the entire dataset. Statistical significance was determined using limma ^32^. A fold-change threshold of ±1.5 (log2 transformed) was applied to identify the downregulated and upregulated genes.

### RiboMethSeq and analysis

The Illumina-based RiboMethSeq protocol was used to measure the levels of all known Nm sites in human rRNA as previously described ^33–35^. Briefly, 100-150 ng of total RNA from MDA-MB-231 wild-type, hTERT-HME1 wild-type, MDA-MB-231 control, and FBL KD1 cells were subjected to random alkaline fragmentation, and, after two-step end repair (3’-dephosphorylation and 5’-phosphorylation), fragments were used as input material for the NEBNext Small RNA kit for Illumina. Library preparation included ligating adapters to the 3’ and 5’ ends, and RT and PCR amplification and barcoding. Barcoded libraries were sequenced using Illumina NextSeq2000, with a target coverage of 10 million raw reads per sample. Bioinformatics treatment included trimming the sequencing adapter, alignment of reads to the human rRNA reference, and counting the reads’ extremities (5’ and 3’). The QuantScore (ScoreC2 or MethScore2 in previous publications) was calculated for the +/-2 nucleotide interval. QuantScore represents the protection of the phosphodiester bond against nucleolytic cleavage upon alkaline fragmentation, and thus linearly correlates with the methylation level, with a maximum level of 1, indicating that all the rRNAs are fully modified at the given site. When methylation levels are low or absent, QuantScore may turn negative because the residue may be more accessible to nucleophiles and, thus, more extensively cleaved than other residues in the environment. Negative QuantScore values are generally neglected, and indicate very low or zero methylation levels. When the QuantScore values were between 0 and 1, this suggested that the samples displayed heterogeneous methylation levels at a given position. Since it is already known that partial RNA degradation affects the precision of RiboMethSeq quantification, the quality of the RNA was assessed by agarose electrophoresis and Bioanalyzer capillary electrophoresis (Bioanalyzer 2200 RNA Pico Chip).

### snoRNA expression analysis

Reads from RiboMethSeq were quality-checked using FASTQC (v0.11.4) ^36^, and TrimGalore (v0.6.0) ^37^ was used for adapter removal and quality trimming. The reads were aligned to the human genome (GRCh38.v38) and quantified using STAR (v2.7.0f) ^38^. Gene counts were pre-filtered, ribosomal genes were removed, and differential gene expression analysis between control and FBL KD cells was then computed using the Deseq2 (v.1.26.0) R package ^27^. Significant genes were retrieved using a threshold of 0.05, the Benjamini-Hochberg adjusted *P* value, and a minimum FC of 1.5 (Log_2_-transformed). Only C/D box snoRNAs were retrieved for further analysis.

### Seahorse cellular stress assays

To evaluate mitochondrial and glycolytic functions, the Seahorse XF96 Cell MitoStress Test and Seahorse XF96 Glycolysis Stress Test (Agilent Technologies, Santa Clara, CA) were performed according to the manufacturer’s protocol. Briefly, control and FBL KD MDA-MB-231 cells were seeded at a density of 2 × 10^4^ cells/well in a Seahorse XF96 Cell Culture Microplate coated with poly-d-lysine (50 μg/ml, Sigma-Aldrich). Cells were cultured overnight at 37 °C in a humidified chamber. After 24 h, cells were washed three times with low-buffered pH 7.4 DMEM (Sigma-Aldrich) supplemented with 2 mM glutamine (Thermo Fisher Scientific^TM^) for GlycoStress assays and with the addition of 10 mM glucose (Thermo Fisher Scientific^TM^) and 1 mM pyruvate (Thermo Fisher Scientific^TM^) for MitoStress assays. The microplate was incubated at 37 °C in a CO_2_-free incubator for 1 h. Compounds were prepared in assay media and added to the injection ports of the assay plate. For MitoStress assays, 1 μM oligomycin (Sigma-Aldrich), 2 μM carbonyl cyanide 4-(trifluoromethoxy) phenyl-hydrazone (FCCP, Sigma-Aldrich), 0.5 μM rotenone (Sigma-Aldrich), and 0.5 μM antimycin A (Sigma-Aldrich) were used. For the GlycoStress assays, 10 mM glucose, 1 μM oligomycin, and 50 mM 2-deoxy glucose (Sigma-Aldrich) were used. After the assay, the cells were stained with 5 μg/ml Hoechst 33342 (Thermo Fisher Scientific^TM^), and fluorescence was measured at 405/450 nm (excitation/emission) using a plate reader (Synergy 2, BioTek). Data were exported from the Seahorse XF96 Extracellular Flux Analyzer into the Seahorse XF Report Generator software. The oxygen consumption rate (OCR) and extracellular acidification rate (ECAR) were normalized to the total cell number, as determined by Hoechst staining, and reported as pmoles/min/cell number for OCR and mpH/min/cell number for ECAR.

### MitoTracker staining

MitoTracker^TM^ Green FM (Invitrogen^TM^) was used to label the mitochondria in control and FBL-KD cells. Briefly, cells were seeded on coverslips coated with 50µM/ml Poly-L-lysine hydrobromide (Sigma-Aldrich) and incubated overnight in a 5% CO_2_ atmosphere at 37 °C. The following day, the cells were incubated with MitoTracker^TM^ Green FM diluted in DMEM without serum at a final concentration of 30 nM for 30 min. Thereafter, media containing Hoechst 33342 (5 μg/ml, Thermo Fisher Scientific^TM^) was added to each well, and the plate was incubated for an additional 10 min. Cells were fixed using a solution of 4% formaldehyde in 1X PBS for 15 min at RT. The coverslips were washed three times with 1X PBS and mounted using SlowFade™ Glass Soft-set Antifade Mountant (Invitrogen^TM^). Images were acquired using a Leica confocal microscope at 63X magnification. Post-acquisition, the intensity of images was adjusted using LAS X (Leica Application Suite X) software to standardize the fluorescence signal intensity across samples for better visualization.

### Protein Synthesis

Global protein synthesis was measured using the Global Protein Synthesis Assay Kit (ab235634, Abcam) according to the manufacturer’s protocol for immunofluorescence with slight modifications. After selection with hygromycin, control and FBL-KD cells were seeded on coverslips in complete media. The following day, the cells were incubated with 100 μl of 1X protein-label solution containing O-propargyl puromycin (OP-puro) for 2 h. A separate control group was incubated with either OP-puro or CHX. Incubation for 15 min with Fixative Solution 1X was used to fix the cells on the coverslips. After that, the cells were permeabilized by incubation with 1X Permeabilization Buffer for 10 min. After permeabilization, 100 μl of the Reaction Cocktail composed of 1X PBS, Copper Reagent, Fluorescent Azide, and Reducing agent was added to each coverslip and incubated for 30 min. Nuclei were stained with Hoechst 33342 (5 μg/ml; Thermo Fisher Scientific^TM^) for 5 min. Finally, the coverslips were washed with 200 μl of 1X PBS and mounted on slides. Images were acquired using a Leica confocal microscope at 63X magnification. Post-acquisition, the intensity of images was adjusted in LAS X (Leica Application Suite X) software to standardize the fluorescence signal intensity across samples for visualization and analysis. The modifications were applied uniformly to all images within the experimental group. The fluorescence intensity was quantified using the threshold function in the Fiji (ImageJ) software. The nuclei were counted using a counting tool. Five images were analyzed for each condition. The results were averaged across three biological replicates.

### Gene Ontology analysis

Gene ontology (GO) analysis for the enrichment of biological processes was performed using the web tool The Database for Annotation, Visualization, and Integrated Discovery (DAVID) (http://david.abcc.ncifcrf.gov/).

### Ethics approval and consent to participate

Patients were recruited through the Sweden Cancerome Analysis Network - Breast (SCAN-B) study (ClinicalTrials.gov ID NCT0230609) ^39^, which received approval from the Regional Ethical Review Board in Lund, Sweden (Registration numbers: 2009/658, 2010/383, 2012/58, 2016/742, 2018/267, and 2019/01252). Written informed consent was obtained from all participants prior to enrollment. To ensure patient confidentiality, all clinical samples were coded. All analyses were conducted in compliance with ethical regulations and patient consent.

### SCAN-B RNA-seq data accessibility

The Swedish Cancerome Analysis Network - Breast (SCAN-B) dataset contained RNA-seq data from 7,743 breast cancer patients ^39^. A dataset of 66 control samples derived from normal breast tissue was also included. These control samples were obtained from healthy women undergoing mammoplasty surgery ^40^.

Clinical chart reviews were conducted to extract data on tumor characteristics, patient demographics, treatment details, and follow-up outcomes, including relapses and mortality. Patients who were alive and relapse-free at the time of data retrieval were censored.

### Differential expression analysis of SCAN-B data

Differential expression (DE) analysis was performed for all pairwise comparisons of breast cancer subtypes according to the SSP classification and for breast cancer subtypes versus normal control samples employing the DESeq2 R package (v1.40.2) ^27^. Technical replicates of SCANB samples were collapsed according to the sample-to-merged associations provided in the supplementary data table ^41^. Genes with fewer than 10 total counts were excluded from the DE analysis. All breast cancer subtypes were compared against the negative control (healthy breast tissue) in the DE analysis. DESeq2’s median of ratios method was employed to normalize gene counts, accounting for differences in sequencing depth and library size between samples. A threshold of adj *P* value < 0.05 and |Log_2_(FC)| > 0.85 was selected to define significantly up- and down-regulated genes. For the mean comparison between different conditions (subtypes, Ki67 level, and ROR), the two-tailed t-test for independent samples was used.

### Overall Survival and disease-free analysis of SCAN-B dataset

Evaluation of the prognostic effect of genes in breast cancer was carried out according to the overall survival (OS) and disease-free (DF) time and events annotated in the SCAN-B supplementary table containing the sample metadata ^41^. OS time (in days) and death status, as well as DF time and relapse status, were employed to construct Kaplan-Meier models using the survival R package (v0.5.0) ^42^, with OS/DF curves generated via survminer (v0.5.0) ^43^. H-score values were stratified into high (H-score > q75) and low (H-score ≤ q25) categories based on the 25^th^ and 75^th^ percentiles. The multiple test correction of log-rank test *P* values was performed using the Benjamini-Hochberg (BH) method.

### Tissue microarrays

The patient cohort for the construction of the tissue microarray (TMA) has been previously described ^44^. TNBC status was determined by immunohistochemistry (IHC) staining for ER, PR, and HER2. Following Swedish National Guidelines, tumors were classified as triple-negative if ER and PR staining was positive in fewer than 10% of tumor cells and HER2 was either scored 0/1+ by IHC or deemed non-amplified by *in situ* hybridization (ISH) if the IHC score was 2+.

As part of the SCAN-B project (NCT02306096), a TMA was constructed, comprising TNBC samples (two 1 mm cores per tumor) from patients diagnosed between September 2010 and December 2014 in the Skåne healthcare region, Sweden. Hematoxylin and eosin (H&E) staining was initially performed to confirm sufficient tumor content (>20% tumor cellularity).

For IHC staining of the TMAs all detection reagents and instrumentation were obtained from Ventana Medical Systems. The TMAs were stained with primary antibodies against FBL (sc-166021; dilution 1:400), RPS28 (14796-1-AP; dilution 1:400), IRAK1 (66653-1-ig; dilution 1:250), and TMSB10 (sc-514309; dilution 1:200) using the automated BenchMark ULTRA platform for 32 min. Heat-induced epitope retrieval was performed using ULTRA Cell Conditioning 1 at 95°C for 36 min in the case of FBL, IRAK1, and TMSB10 or ULTRA Cell Conditioning 2 at 91°C for 24 min in the case of RPS28. Primary antibodies were detected and visualized with Ultraview Universal DAB Detection Kit. Hematoxylin was used as counterstain.

### H-score

The histochemical score (H-score) was determined by manually assessing the intensity of staining for each core in the TMA (0 = no staining, 1+ = weak, 2+ = moderate, and 3+ = strong) and multiplying it by the percentage of stained cells (0–100%), resulting in a score ranging from 0 to 300. Two cores were analyzed per patient, and the final H-score was determined as the average of the two cores. All pathology assessments were conducted blinded to the pathological and molecular classifications of the samples.

### Statistics

Statistical analyses for experimental data were performed using GraphPad Prism software (version 10.0.0 and higher) for Windows (GraphPad Software, La Jolla, California, USA). The significance was determined using different statistical tests as specified. In the case of multiple comparisons, specific correction methods were applied as specified. Probability (*P*) values or adjusted *P* values (adj *P*) of \**P* < 0.05, \*\**P* < 0.01, \*\*\**P* <0.001, \*\*\*\**P* < 0.0001 were considered statistically significant. Statistical analysis of high-throughput and patient-related data was performed in R Studio. For all patient-related analyses, the unpaired two-tailed t-test corrected with the BH method for multiple comparisons, * adj *P* < 0.05, ** adj *P* < 0.01, *** adj *P* < 0.001, **** adj *P* < 0.0001, ns adj *P* > 0.05.

## Results

### FBL sustains cell growth and proliferation in TNBC

To explore the function of FBL in TNBC cells, we first assessed the protein and mRNA expression levels in the TNBC cell line MDA-MB-231 and the normal mammary epithelial cell line, hTERT-HME1. Our analysis revealed increased expression of FBL at both the protein and mRNA levels in MDA-MB-231 cells compared with that in the normal cell line (**Figures 1A** and **1B**). Similarly, analysis of the Sweden Cancerome Analysis Network-Breast (SCAN-B) breast cancer cohort showed *FBL* mRNA expression was elevated across all breast cancer subtypes-except for the luminal B subtype-when compared to normal tissue (**Figure 1C**). Moreover, FBL expression correlated with Ki67, a well-established proliferation marker (**Figure 1D**).

**Figure 1.**
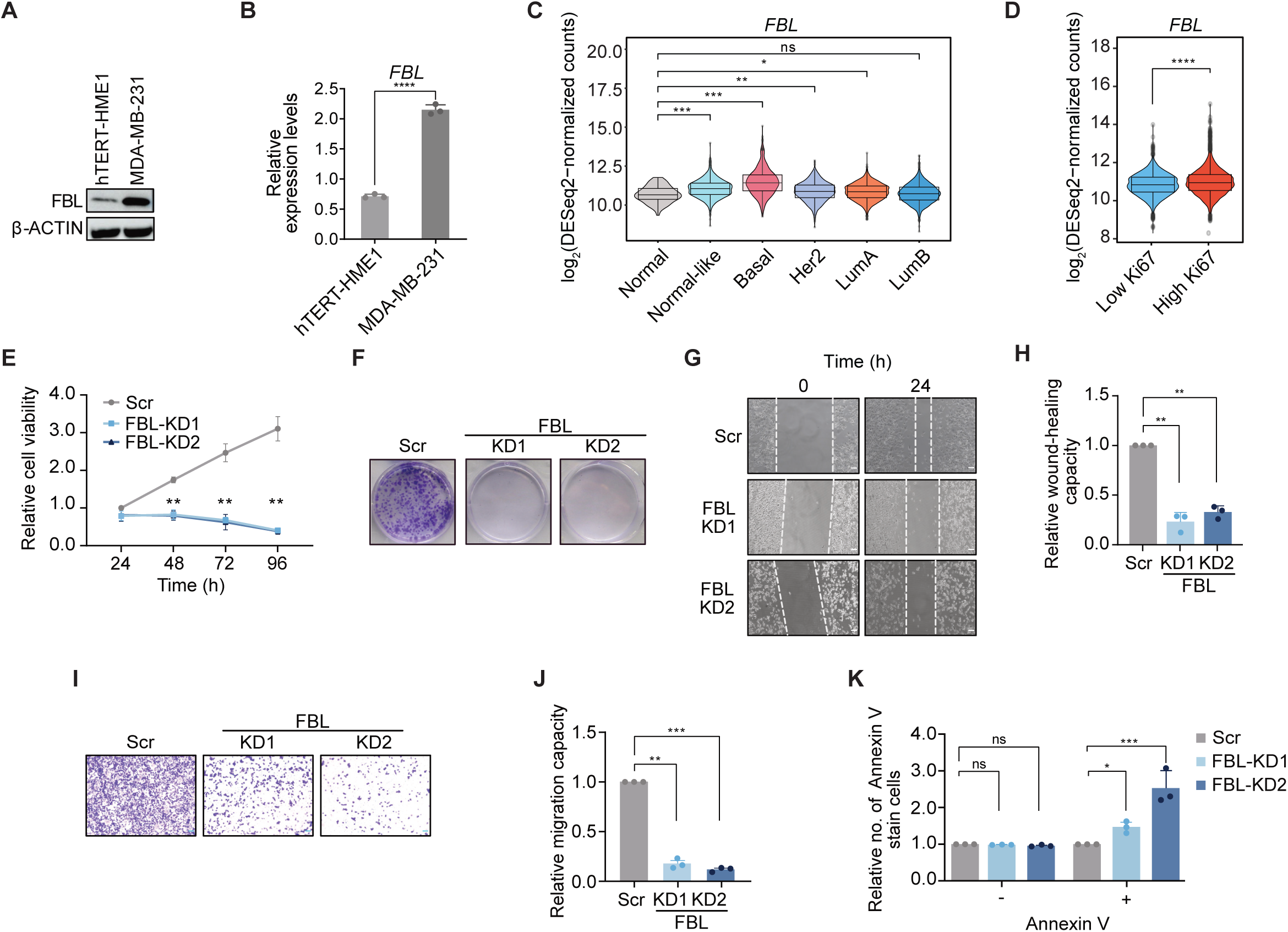
FBL is essential for cell growth and migration in MDA-MB-231 cells. (**A**) Representative Western Blot depicting FBL expression in hTERT-HME1 and MDA-MB-231 cell lines. β-ACTIN serves as a loading control. (**B**) RT-qPCR analysis of *FBL* in hTERT-HME1 and MDA-MB-231 cell lines. The mRNA expression levels were normalized to *GAPDH*. (**C**) *FBL* expression across breast cancer subtypes in the SCAN-B cohort. (**D**) Violin plot illustrating *FBL* expression in SCAN-B patient samples categorized by low and high Ki67 levels. (**E**) MTT assay of control and FBL-depleted (KD1 and KD2) cells over 4 days (24, 48, 72, and 96 h). (**F**) Representative images of the crystal violet assay of control and FBL-depleted (KD1 and KD2) cells after 10 days in culture. (**G**) Representative images from the wound-healing assay showing the wound-healing capacity of control and FBL-depleted (KD1 and KD2) cells upon scratch generation at 0 and 24 h. Scale bar: 100 µm. (**H**) Column plot showing quantification of the wound-healing capacity of FBL-depleted (KD1 and KD2) cells relative to control based on the wound-healing assay. (**I**) Representative images showing control, FBL-KD1, and KD2 cell migration through the trans-well. Scale bar: 100 µm. (**J**) Column plot showing quantification of the migration capacity of FBL-depleted (KD1 and KD2) cells relative to control based on the trans-well assay. (**K**) Apoptosis analysis for control and FBL-depleted (KD1 and KD2) cells plotted as Annexin V negative (-) and Annexin V (+) cells. Statistical analysis: All experimental data are presented as mean ± SD, n = 3. (**B**) Unpaired two-tailed t-test, * *P* < 0.05, ** *P* < 0.01, **** *P* < 0.0001. (**C** and **D**) Unpaired two-tailed t-test corrected with the FDR method for multiple comparison, * adj *P* < 0.05, ** adj *P* < 0.01, *** adj *P* < 0.001, **** adj *P* < 0.0001, ns adj *P* > 0.05. (**E** and **K**) Two-way ANOVA with Dunnett’s multiple comparisons test, * adj *P* < 0.05, *** adj *P* < 0.001. (**H** and **J**) Paired two-tailed t-test, ** *P* < 0.0001, *** *P* < 0.0001.

To further explore FBL function, we employed a loss-of-function approach with two distinct shRNAs (KD1 and KD2) targeting *FBL* mRNA. KD efficiency was assessed by western blotting and RT-qPCR, demonstrating a nearly complete depletion of FBL (**Figures S1A** and **S1B**). The MTT assay showed a significant reduction of approximately 88% in cell proliferation upon FBL silencing over a 96 h period (**Figure 1E**). To assess whether the cells were viable after puromycin selection, we stained them with calcein AM, which showed no significant difference in viability between Scr and KD cells (**Figure S1C** and **S1D**). This suggests that although cell growth was reduced, the cells did not die and retained membrane integrity. In addition, crystal violet staining revealed that the clonogenic capacity of FBL-KD cells was also impaired, as these cells could not form colonies (**Figure 1F**). To evaluate the migration capacity, we conducted wound-healing and transwell assays, which demonstrated a greater than 65% reduction in wound-healing capacity and an over 85% decrease in migratory capacity of FBL KD cells compared to controls (**Figure 1G-J**). Apoptosis analysis revealed a significant increase in Annexin V-positive cells following FBL silencing (**Figures 1K** and **S1E**). Altogether, these data suggest that FBL influences breast cancer tumorigenesis, as its depletion results in decreased proliferation, impaired colony formation, and reduced migration of TNBC cells.

### FBL controls translation and mitochondrial function

Given the role of FBL in ribosome biogenesis ^45^, we assessed its impact on translation in TNBC cells using polysome profiling in control and FBL-KD cells. FBL-depleted cells exhibited an overall decrease in ribosomal subunits, monosomes, and polysomes compared with control cells (**Figure 2A**). Furthermore, FBL depletion led to the accumulation of stalled pre-initiation complexes, specifically halfmers, composed of an 80S ribosome and a single 43S pre-initiation complex (**Figure 2A**). These halfmers could arise from ribosome biogenesis defects caused by underproduction of 60S subunits or impaired subunit joining ^46,47^. Consistent with this, FBL silencing resulted in reduced levels of 18S and 28S rRNAs (**Figure S2A**). Overall, these findings suggest that FBL depletion drastically reduced ribosome availability, thereby impairing global translation. Supporting this, a *de novo* protein synthesis assay based on OP-Puro incorporation confirmed a marked decrease in protein synthesis upon FBL silencing (**Figures 2B, 2C,** and **S2B**).

**Figure 2.**
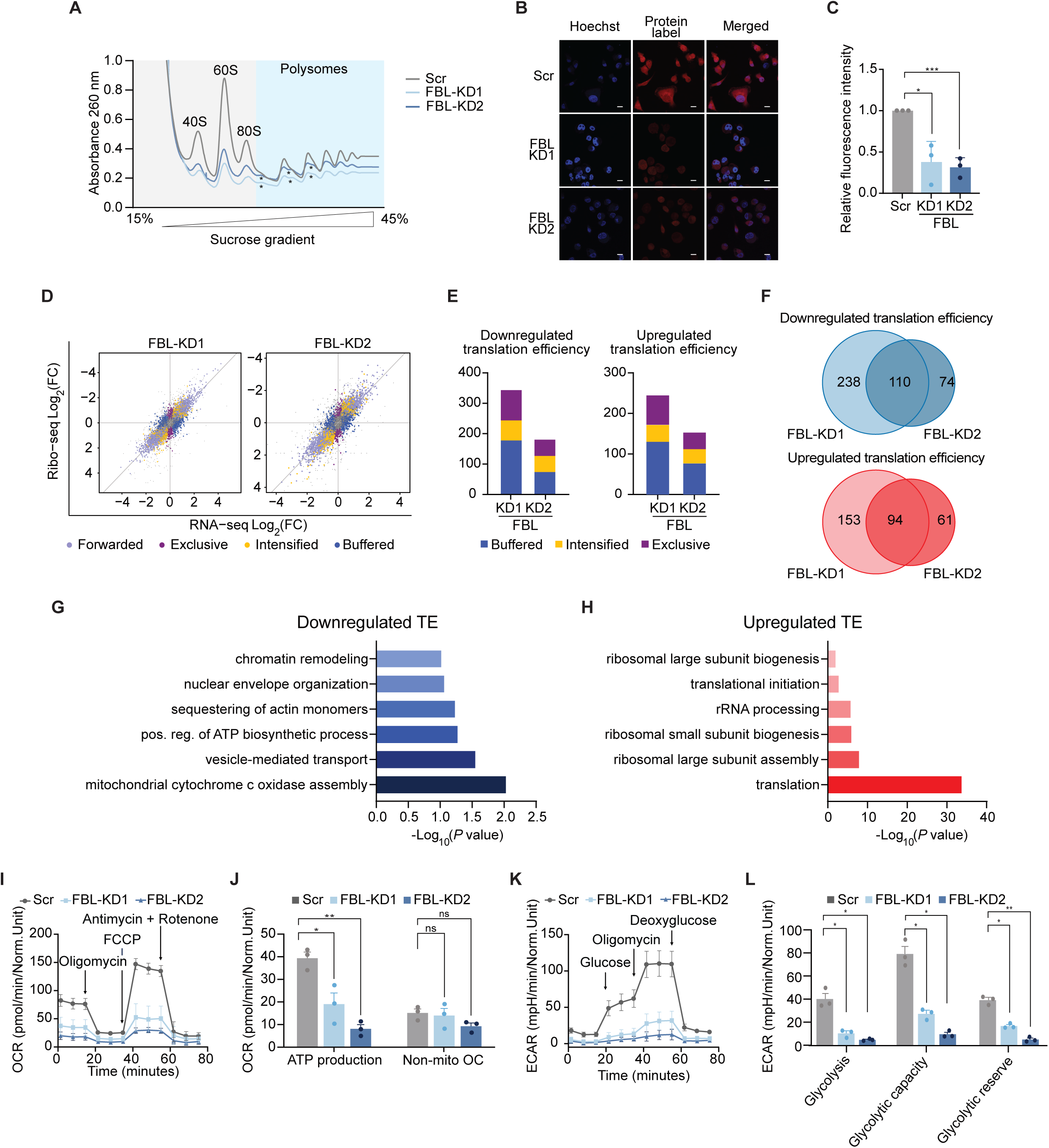
Silencing of FBL induces subtle changes in translation, influencing metabolic parameters. (**A**) Polysome profiling for control and FBL-depleted (KD1 and KD2) cells in a 15-45% sucrose gradient. Peaks for 40S, 60S, 80S, and polysomes are highlighted and correspond to absorbance measurements at 260 nm. Halfmers are marked with *. (**B**) Representative images of nascent proteins labeled in control and FBL-depleted (KD1 and KD2) cells. Hoechst 33342 was used to mark the nuclei. The protein label solution was used to label *de novo* peptides. Scale bar: 10 µm. (**C**) Quantification of immunofluorescence staining for protein label signal. (**D**) Scatter plot showing the classification of genes according to the Fold changes (FC) of and ribosome-protected fragments (RPF), mRNA, and translation efficiency (TE). The Y-axis shows Ribo-Seq Log_2_(FC), X-axis shows RNA-Seq Log_2_(FC). The genes are classified as buffered (dark blue), intensified (yellow), exclusive (dark purple), and forwarded (light purple). (**E**) Column plot representing buffered, intensified, and exclusive transcripts with downregulated (left) and upregulated (right) translation efficiency in KD1 and KD2. (**F**) Venn diagram showing the overlap of genes with downregulated (blue) and upregulated (red) translation efficiency between the two FBL KDs. (**G** and **H**) GO analysis of biological processes associated with genes with (**G**) downregulated or (**H**) upregulated translation efficiency upon FBL depletion. (**I**) Oxygen consumption rate (OCR) in control and FBL-depleted (KD1 and KD2) cells. Traces display OCR during basal conditions and upon injection with cellular respiration-modulating compounds (oligomycin, FCCP, antimycin A with rotenone) at the indicated time points. (**J**) Mitochondrial respiration and function parameters in control and FBL-depleted (KD1 and KD2) cells based on the OCR: ATP production and non-mitochondrial oxygen consumption (OC). (**K**) Extracellular acidification rates (ECAR) in control and FBL-depleted (KD1 and KD2) cells. Traces display ECAR during basal conditions with no glucose and upon injection of glucose, oligomycin, and 2-deoxy-glucose at the indicated time points. (**L**) Glycolysis parameters in control and FBL-depleted (KD1 and KD2) cells were calculated based on ECAR: non-glycolytic acidification, glycolysis, glycolytic capacity, and glycolytic reserve. Statistical analysis: All data are presented as mean ± SD, n = 3. (**C**) Unpaired two-tailed t-test, * *P* < 0.05, *** *P* < 0.001. (**J** and **L**) Two-way ANOVA with Dunnett’s multiple comparisons test. * adj *P* < 0.05, ** adj *P* < 0.01. The cut-off applied to the Ribo-Seq data analysis was Log_2_(FC) ± 0.585, adj *P <* 0.05.

To investigate whether FBL modulates the translation efficiency of specific mRNAs, we performed Ribo-seq on the control and FBL-depleted cells. Quality control analysis of the footprints confirmed alignments in the CDS regions and periodicity of the 5’ ends of Ribo-Seq reads for both controls and FBL-KD cells (**Figure S2C**-**F**). RNA-Seq analysis of the input samples identified 4,783 differentially transcribed genes (DTGs) in KD1 and 3,701 in KD2 following FBL silencing (**Figures S3A** and **S3B; Table S2**). Among these, 2,469 and 2,013 genes were downregulated, whereas 2,314 and 1,688 were upregulated in FBL-KD1 and FBL-KD2, respectively (**Figures S3C-E**). Gene ontology (GO) analysis of commonly downregulated genes revealed categories linked to fundamental mechanisms governing cell growth maintenance, including cell division, DNA replication, and mitotic spindle organization (**Figure S3F**). Conversely, commonly upregulated genes were associated with cell adhesion, cell migration, collagen fibril, and extracellular matrix organization (**Figure S3G**).

In contrast, the Ribo-seq data revealed a lower number of genes with significant deregulation at the translational level (differentially translated genes (DTEGs)), with 623 and 377 genes for KD1 and KD2, respectively (**Table S3**). Furthermore, we categorized the DTEGs based on their differential regulation as follows: i) transcriptionally forwarded genes, which are regulated at the transcriptional level and exhibit significant changes in both mRNA and ribosome-protected fragments (RPF) without a notable change in translation efficiency; ii) translationally exclusive genes, representing genes that show significant changes in RPF and translation efficiency, but no corresponding changes in mRNA levels; iii) translationally intensified genes, exhibiting significant, aligned changes in mRNA, RPF, and translation efficiency; and iv) translationally buffered genes, characterized by significant changes in mRNA, RPF, and translation efficiency, but with transcriptional changes counteracting the alterations in translation efficiency (**Figure 2D**) ^28^. Among the genes with significantly downregulated translation efficiency in FBL-KD1, 181 were categorized as buffered, 66 as intensified, and 100 as exclusive. In FBL-KD2, 77 were buffered, 50 were intensified, and 57 were excluded. For genes with a significant increase in translation efficiency, 132 were buffered, 42 intensified, and 73 were exclusive to FBL-KD1, whereas FBL-KD2 included 79 buffered, 35 intensified, and 41 exclusive transcripts (**Figure 2E**). Overlap analysis between both knockdowns identified 110 transcripts with downregulated translation and 94 transcripts with upregulated translation (**Figure 2F**). GO analysis of transcripts with downregulated translation efficiency revealed a broad range of processes, including mitochondrial cytochrome C oxidase assembly, vesicle-mediated transport, positive regulation of ATP biosynthesis, and actin monomer sequestration (**Figure 2G**). In contrast, transcripts with upregulated translation efficiency were enriched in the categories related to translation, rRNA processing, and ribosome biogenesis. This suggests a potential compensatory mechanism in which cells may redistribute available ribosomes to enhance the production of ribosome biogenesis factors (**Figure 2H**).

To investigate the potential effects of gene expression changes on cellular bioenergetics, we characterized the metabolic profiles of FBL-KD cells by measuring the oxygen consumption rate (OCR) and extracellular matrix acidification rate (ECAR) using Seahorse. Notably, FBL KD cells exhibited significantly reduced OCR under both basal and maximal (FCCP-stimulated) respiration conditions compared to control cells (**Figure 2I**). Correspondingly, ATP production (decrease in OCR upon injection of the ATP synthase inhibitor oligomycin) was also significantly reduced, indicating that FBL-depleted cells were under energetic stress (**Figure 2J**). We confirmed the presence of intact mitochondria in FBL-KD cells using Mitotracker^TM^ Green staining, which was higher in KD cells (**Figure S3H**). Non-mitochondrial oxygen consumption was unaffected by KD (**Figure 2J**), as was coupling efficiency and proton leak (**Figures S3I** and **S3J**). All other MitoStress parameters analyzed were significantly reduced in FBL-depleted cells, supporting a dysfunctional mitochondrial phenotype (**Figure S3K**). The glycolysis Stress assay revealed impaired ECAR in KD cells compared to control cells (**Figure 2K**). Hence, glycolysis parameters were also significantly decreased in KD cells (**Figures 2L** and **S3L**). Collectively, these findings suggest that FBL depletion affects translation, which has an effect on both mitochondrial and glycolytic functions, uncovering potential vulnerabilities in FBL-deficient TNBC cells.

### FBL maintains global protein synthesis and translation of proto-oncogenes

To assess changes in protein abundance over time, we conducted mass spectrometry-based proteomics on control and FBL-depleted cells after 48 and 96 h of puromycin selection. In total, 7,668 proteins were identified in the samples (**Table S4**). At 96 h, we observed a global decrease in protein abundance relative to the 48 h time point and the control cells (**Figure 3A**). Overlapping of the significantly downregulated proteins across both time points revealed 4,019 common proteins (**Figure 3B**). GO analysis indicated that these proteins were enriched in biological processes, such as translation, cell division, cell cycle, and chromatin remodeling (**Figure 3C**). Additionally, the enrichment of protein folding, transport, and phosphorylation categories suggested that FBL depletion may cause broader functional disruptions (**Figure 3C**).

**Figure 3.**
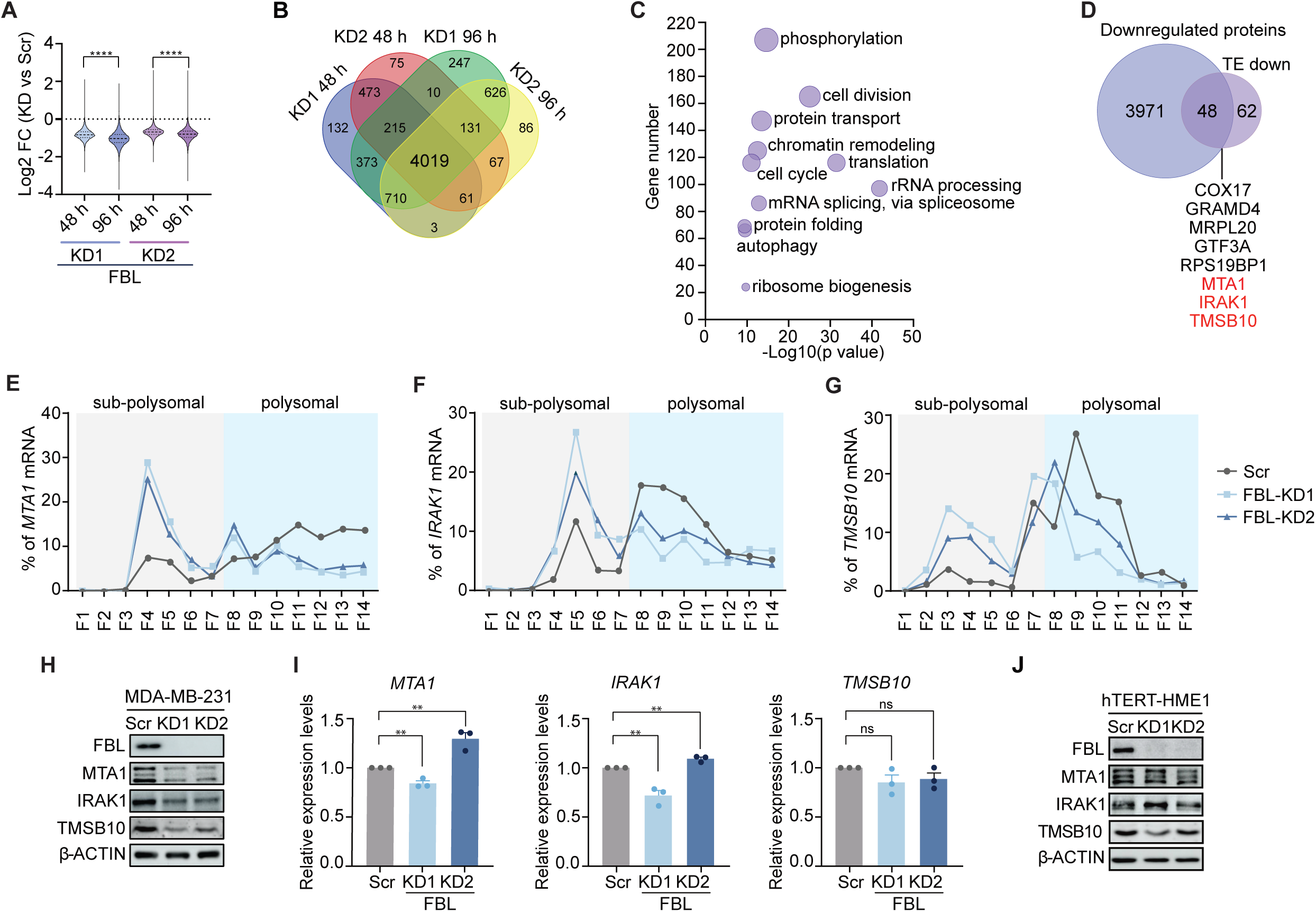
FBL regulates the translation of proto-oncogenes. (**A**) Violin plot illustrating the distribution of proteins detected in total proteomics based on the Log_2_(FC) compared to control for both KDs after 48 and 96 h of FBL depletion. (**B**) Venn diagram displaying the overlap of downregulated proteins at 48 h and 96 h in both FBL KDs. (**C**) Common enriched biological processes associated with downregulated proteins upon FBL KDs. (**D**) Venn diagram of the overlap between the commonly downregulated proteins, and the transcripts with downregulated translation efficiency. Highlighted in red are proteins with an already described role in breast cancer. (**E**-**G**) mRNA percentage distribution among the 14 fractions collected by polysome profiling for (**E**) *MTA1*, (**F**) *IRAK1*, and (**G**) *TMSB10* in control and FBL-depleted cells. Ribosomal subunits and 80S fractions are highlighted in light grey, and polysome fractions are highlighted in blue. (**H**) Representative Western Blot images displaying the expression of FBL, MTA1, IRAK1, and TMSB10 in control and FBL-depleted (KD1 and KD2) MDA-MB-231 cells. β-ACTIN serves as a loading control. (**I**) RT-qPCR analysis of *MTA1*, *IRAK1,* and *TMSB10* levels from control and FBL-depleted (KD1 and KD2) MDA-MB-231 cells. mRNA expression levels were normalized to *GAPDH* and are presented relative to the control. (**J**) Representative Western Blot images displaying the expression of FBL, MTA1, IRAK1, and TMSB10 in control and FBL-depleted (KD1 and KD2) hTERT-HME1 cells. β-ACTIN serves as a loading control. Statistical analysis: All data are presented as the mean ± SD, n = 3. (**A**) Welch’s two-tailed t-test; **** *P* < 0.0001.(**I**) Unpaired two-tailed t-test, * *P* < 0.05, ** *P* < 0.01, **** *P* < 0.0001. The cut-off applied to proteomic data analysis was Log_2_(FC) ± 0.585, adj *P <* 0.05.

Intersecting these proteins with transcripts exhibiting reduced translation efficiency highlighted 48 proteins (**Figure 3D**). Among these, several are associated with mitochondrial functions (e.g., COX17, GRAMD4, and MRPL20) ^48–50^, others with rRNA transcription and processing (e.g., GTF3A and RPS19BP1) ^51,52^, and some processes linked to breast cancer progression (e.g., MTA1, IRAK1, and TMSB10) ^53–58^ (**Figure 3D** and **S4A-I**). To investigate FBL-mediated translation regulation of the proto-oncogenes *MTA1*, *IRAK1*, and *TMSB10*, we conducted polysome profiling followed by RT-qPCR in control and FBL KD cells, analyzing mRNA from different fractions, including those containing unbound mRNAs, ribosomal subunits, and mRNAs bound by varying numbers of ribosomes. The results revealed that in FBL KD cells, *MTA1*, *IRAK1,* and *TMSB10* mRNAs accumulated in the subpolysomal fractions, indicating disrupted translation. In contrast, the control cells exhibited a higher percentage of these transcripts in the polysome-containing fractions (**Figures 3E-G** and **S4J-O**). Furthermore, the distribution of *GAPDH* mRNA across fractions demonstrated the specificity of the observed effect, as *GAPDH* mRNA did not accumulate in the non-polysome fractions in FBL-KD cells (**Figures S4P-R**).

Next, we assessed protein levels in MDA-MB-231 cells and observed a marked reduction in all three candidates upon FBL silencing (**Figure 3H**). This reduction in protein levels could not be attributed to changes in steady-state mRNA levels of these proteins (**Figure 3I**). Interestingly, in the normal mammary epithelial cell line hTERT-HME1, FBL silencing had a different impact on the protein expression of MTA1, IRAK1, and TMSB10. TMSB10 showed a mild reduction, whereas MTA1 and IRAK1 protein levels remained unaffected (**Figure 3J**). Altogether, our findings indicate that FBL-mediated regulation of MTA1, IRAK1, and TMSB10 expression occurs at the translational level in TNBC cells, and that this effect is specific to breast cancer cells.

### TNBC cells exhibit a specific Nm signature

Given the main role of FBL in depositing Nm on rRNAs, we employed RiboMethSeq to map Nm sites on rRNA in MDA-MB-231 and hTERT-HME1 cell lines ^35^. Human rRNA has 110 Nm sites distributed as follows: 41 on 18S rRNA, 67 on 28S rRNA, and 2 on 5.8S rRNA. Each methylation site was assigned a QuantScore representing the fraction of methylated molecules at each position. The QuantScore distributions were comparable between the cell lines, with no significant differences (**Figures 4A** and **S5A**). Furthermore, the methylation levels between the two cell lines showed a strong positive correlation (r = 0.9152; **Figure 4B**). In MDA-MB-231 cells, 63 sites were almost fully methylated (QuantScore > 0.95) and 15 were partially methylated (QuantScore < 0.75), compared to 68 fully methylated and 15 partially methylated sites in hTERT-HME1 cells (**Table S5**). Notably, 18S-Cm1440 showed no detectable methylation (QuantScore = 0) and 18S-Gm1447 exhibited near-zero methylation levels in both cell lines (**Figure 4C**).

**Figure 4.**
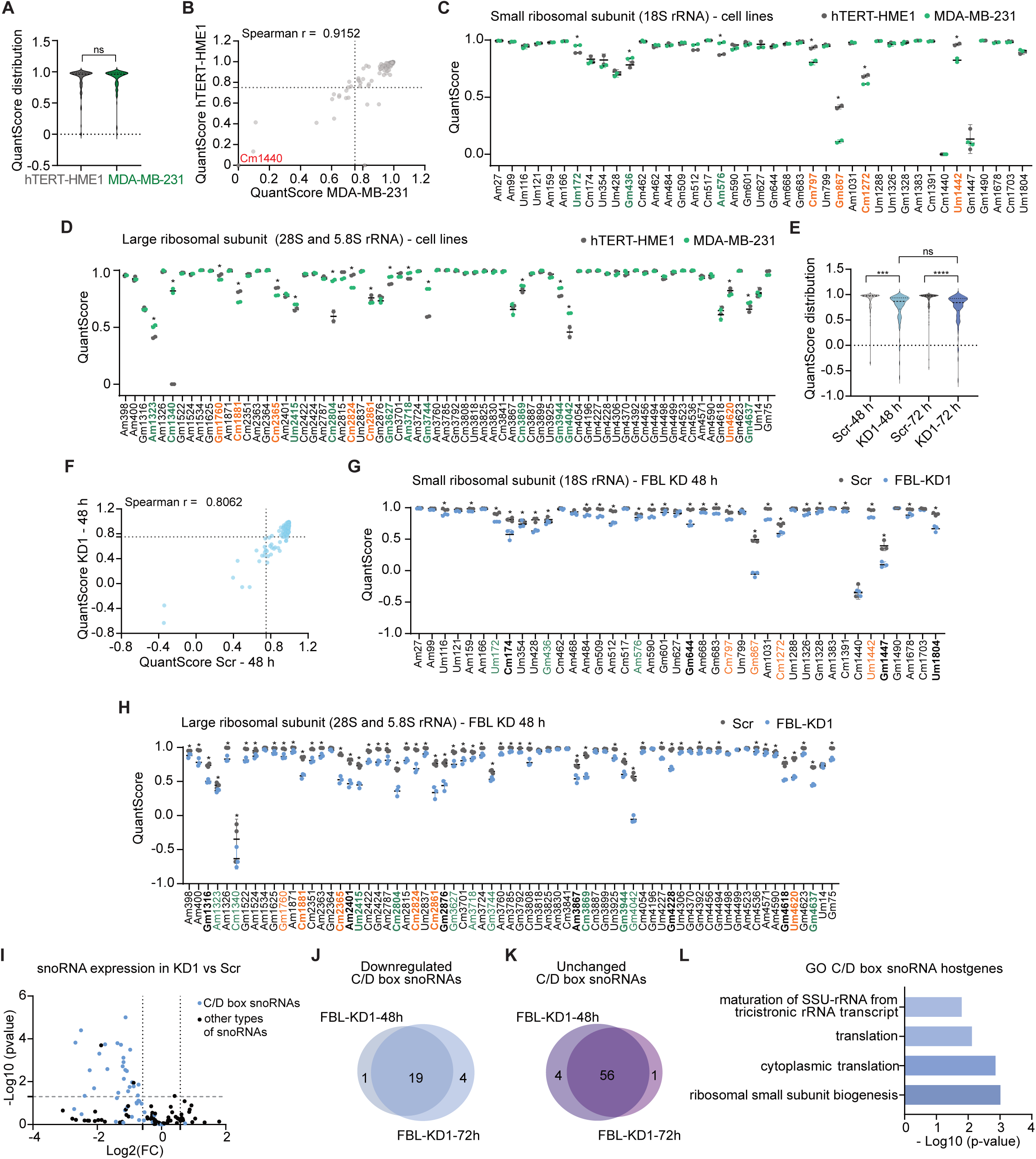
rRNA 2′-O-Methylation profiles and sensitivity to FBL depletion in the TNBC cell line MDA-MB-231. (**A**) Violin plot showing the distribution of QuantScore among hTERT-HME1 and MDA-MB-231 cells. (**B**) Correlation plot between the QuantScore from hTERT-HME1 and MDA-MB-231, with a threshold of 0.75 marked with a dashed line indicating partially methylated sites. (**C** and **D**) Dot plot depicting Nm levels in hTERT-HME1 (grey) and MDA-MB-231 (green) cells within (**C**) the small ribosomal subunit (SSU) containing the 18S rRNA and (**D**) within the large ribosomal subunit (LSU) containing the 28S and 5.8S rRNAs. The X-axis represents modified nucleotide positions, and the Y-axis represents QuantScore for each site. Highlighted in green are sites with higher methylation levels, and in orange are the sites with lower methylation levels in MDA-MB-231 compared to hTERT-HME1. In bold are sites with FC ≥ 1.3 (Scr vs KD1). **E.** Violin plot illustrating the distribution of QuantScore in control (Scr) and FBL-depleted (KD1) cells after 48 and 72 h of FBL depletion. (**F**) Correlation plot between the QuantScore from control and FBL-depleted (KD1) cells, with a threshold of 0.75 indicating partially methylated sites. (**G** and **H**). Dot plot depicting the Nm levels in Scr (grey) and FBL-depleted (KD1) cells (blue) after 48 h of FBL depletion within (**G**) SSU containing the 18S rRNA and within (**H**) LSU containing the 28S and 5.8S rRNAs. The X-axis represents modified nucleotide positions, and the Y-axis represents QuantScore for each site. Highlighted in green are sites with higher methylation levels, and in orange are the sites with lower methylation levels in MDA-MB-231 compared to hTERT-HME1. (**I**) Volcano plot depicting the expression of snoRNAs in KD1 compared to control cells at 48 h of depletion. Data analysis was conducted using the DeSeq2 package. Highlighted in blue are the C/D box snoRNAs. (**J** and **K**) Venn Diagram depicting the overlap of C/D snoRNAs (**J**) downregulated and (**K**) unchanged at 48 and 72 h upon FBL depletion. **L.** Enriched biological processes associated with host genes of downregulated C/D box snoRNAs upon FBL KDs at 48 h. Statistical analysis: (**A** and **E**) Welch’s two-tailed t-test, *** *P* < 0.001,**** *P* < 0.0001, ns = non-significant. (**B** and **F**) Spearman correlation, 95% confidence interval, two-tailed *P* value, **** *P* < 0.0001. (**C, D, G,** and **H**) Two-way ANOVA with the two-stage linear step-up procedure of the Benjamini, Krieger, and Yekutieli test for multiple comparisons. For simplicity, significantly different sites are marked with *, regardless of the *P* value. The threshold for *P* value was set to less than 0.01. (**C** and **D**) Data are presented as the mean ± SD, n = 2. (**G** and **H**) Data are presented as the mean ± SD, n = 3.

Compared to hTERT-HME1 cells, MDA-MB-231 cells exhibited significantly higher QuantScores at 14 sites (3 on the 18S rRNA and 11 on the 28S rRNA; highlighted in green) and significantly lower QuantScores at 10 sites (4 on the 18S rRNA and 6 on the 28S rRNA; highlighted in orange) (**Figures 4C, 4D** and **Table 1**). The largest QuantScore difference was observed for 28S-Cm1340, previously identified as a highly variable site ^59^. Methylation of 18S-Am576 was significantly higher in the TNBC cell line than in the normal mammary epithelial cell line and was previously shown to be increased in TNBC cell lines and human samples ^22^. This suggests that methylation at this position may play a role in TNBC tumorigenesis, making cell lines with similar Nm patterns valuable models for further studies.

**Table 1.** Dysregulated Nm sites in the MDA-MB-231 cell line. The sites with increased (left column) and decreased (right column) significant methylation levels in MDA-MB-231 cells compared to hTERT-HME1 cells. The snoRNA responsible for guiding methylation at each specific site is also indicated.

| rRNA | Increased methylation levels<br>MDA-MB-231 vs. hTERT-HME1 |  | Decreased methylation levels<br>MDA-MB-231 vs. hTERT-HME1 |  |
| --- | --- | --- | --- | --- |
|  | Methylated site | Guide snoRNA | Methylated site | Guide snoRNA |
| 18S | Um172 | SNORD45A/B | Cm797 | SNORD138 |
|  | Gm436 | SNORD100 | Gm867 | SNORD98 |
|  | Am576 | SNORD93 | Cm1272 | SNORD66 |
|  |  |  | Um1442 | SNORD61 |
| 28S | Am1323 (1310) | SNORD126 | Gm1760 (1747) | SNORD73A |
|  | Cm1340 (1327) | SNORD104 | Cm1881 (1868) | SNORD48 |
|  | Um2415 (2402) | SNORD143,<br>SNORD144 | Cm2365 (2352) | SNORD24 |
|  | Cm2804 (2791) | SNORD55 | Cm2824 (2811) | SNORD95 |
|  | Gm3627 (3606) | SNORD31 | Cm2861 (2848) | SNORD50A,<br>SNORD50B |
|  | Am3718 (3697) | SNORD37 | Um4620 (4590) | SNORD72 |
|  | Gm3744 (3723) | SNORD87 |  |  |
|  | Cm3869 (3697) | SNORD53 |  |  |
|  | Gm3944 (3923) | SNORD111 |  |  |
|  | Gm4042 (4020) | SNORD102 |  |  |
|  | Gm4637 (4607) | SNORD121A,<br>SNORD121B |  |  |

To understand the effect of FBL depletion on the stability of Nm sites, we performed RiboMethSeq in control and FBL-KD1 cells 48 and 72 h after FBL depletion (**Figures S6A**, **S7A,** and **Table S6**). A comparison of KD1 QuantScores with those of control cells revealed a significant overall reduction in methylation levels at both time points, with no significant changes between 48 and 72 h in KD1 cells (**Figure 4E**). The correlation between the QuantScores in the control and KD1 cells progressively weakened over time, suggesting a nonlinear dynamic of Nm level reduction upon FBL depletion (**Figures 4F** and **S8A**). Of the 110 sites detected, 72 sites showed a significant (*P* < 0.01) decrease at 48 h, with 27 sites on 18S rRNA, 44 sites on 28S, and one on 5.8S rRNA (**Figure 4G** and **4H**). At 72 h, 12 additional sites showed a significant reduction, with four more sites on 18S rRNA and eight on 28S rRNA (**Figures S8B** and **S8C**). Of the 2 Nm sites on 5.8S, Gm75 showed a significant decrease at both time points **(Figures S8B** and **S8C**).

To identify sites particularly sensitive to FBL depletion, we calculated the FC in QuantScores between control and KD1 cells for each site, defining a meaningful difference as a ≥30% change (FC ≥ 1.3). At 48 h, 20 sites met this threshold, with an additional 6 sites surpassing it by 72 h (**Table 2**). Of these 20 sites, 10 were located on the 28S rRNA (Cm1881, Cm2365, Um2415, Cm2804, Cm2824, Cm2861, Cm3869, Gm3944, Um4620, and Gm4637) and were also differentially methylated in MDA-MB-231 cells (**Figure S8D** and **Table 2**). These findings suggest that certain Nm sites are particularly sensitive to fluctuations in FBL expression, potentially serving roles in fine-tuning translation and influencing oncogenic processes in tumorigenesis, as previously proposed ^60^.

**Table 2.**
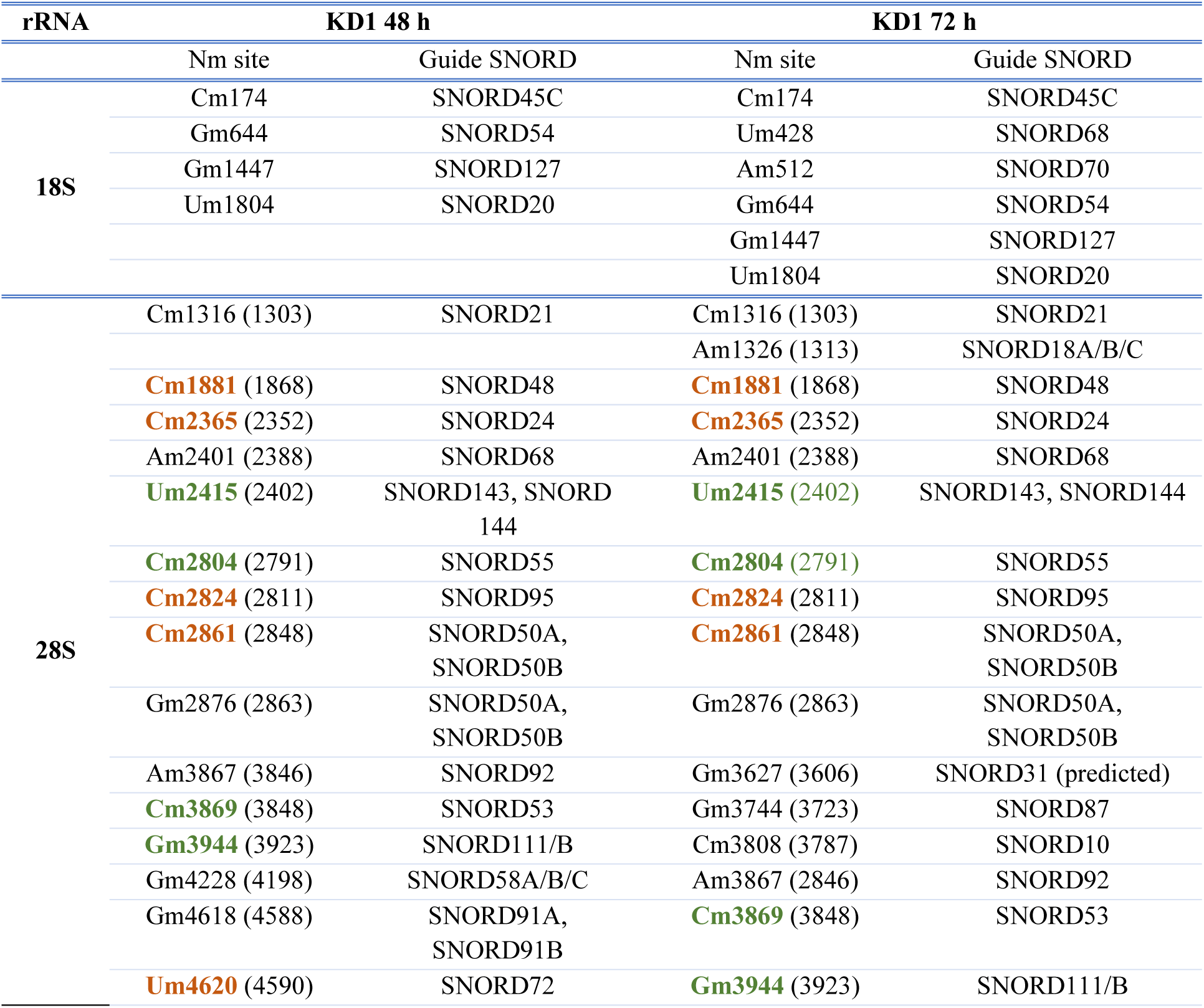
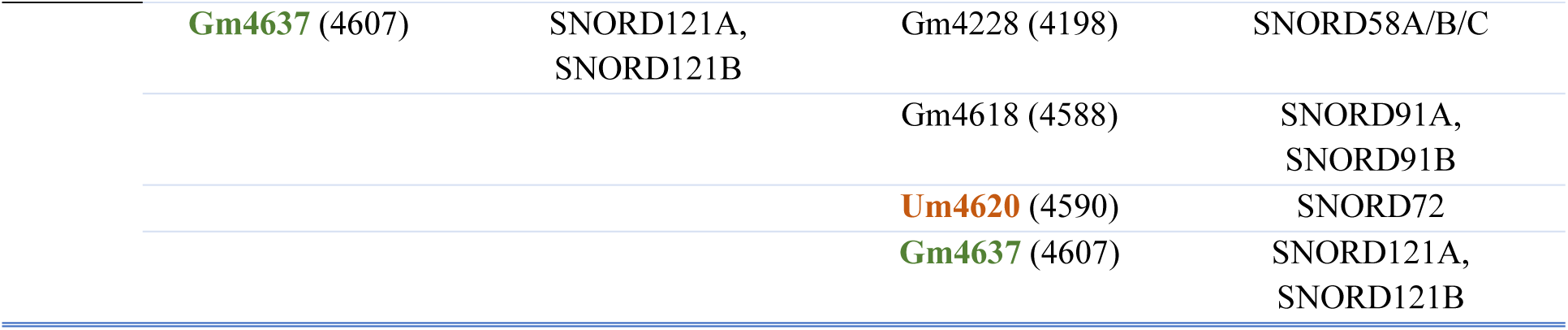
Sensitive sites to FBL depletion. Comparison of QuantScores between control and FBL KD cells identified 20 sites with an FC higher than 1.3 at 48 h and 26 sites at 72 h following FBL depletion. Of these, 10 sites exhibited differential methylation scores in the MDA-MB-231 cell line compared to the normal cell line hTERT-HME1 (highlighted in bold green (higher methylation) and bold orange (lower methylation)).

| rRNA | KD1 48 h |  | KD1 72 h |  |
| --- | --- | --- | --- | --- |
|  | Nm site | Guide SNORD | Nm site | Guide SNORD |
| 18S | Cm174 | SNORD45C | Cm174 | SNORD45C |
|  | Gm644 | SNORD54 | Um428 | SNORD68 |
|  | Gm1447 | SNORD127 | Am512 | SNORD70 |
|  | Um1804 | SNORD20 | Gm644 | SNORD54 |
|  |  |  | Gm1447 | SNORD127 |
|  |  |  | Um1804 | SNORD20 |
| 28S | Cm1316 (1303) | SNORD21 | Cm1316 (1303) | SNORD21 |
|  |  |  | Am1326 (1313) | SNORD18A/B/C |
|  | **Cm1881** (1868) | SNORD48 | **Cm1881** (1868) | SNORD48 |
|  | **Cm2365** (2352) | SNORD24 | **Cm2365** (2352) | SNORD24 |
|  | Am2401 (2388) | SNORD68 | Am2401 (2388) | SNORD68 |
|  | **Um2415** (2402) | SNORD143, SNORD 144 | **Um2415** (2402) | SNORD143, SNORD144 |
|  | **Cm2804** (2791) | SNORD55 | **Cm2804** (2791) | SNORD55 |
|  | **Cm2824** (2811) | SNORD95 | **Cm2824** (2811) | SNORD95 |
|  | **Cm2861** (2848) | SNORD50A, SNORD50B | **Cm2861** (2848) | SNORD50A, SNORD50B |
|  | Gm2876 (2863) | SNORD50A, SNORD50B | Gm2876 (2863) | SNORD50A, SNORD50B |
|  | Am3867 (3846) | SNORD92 | Gm3627 (3606) | SNORD31 (predicted) |
|  | **Cm3869** (3848) | SNORD53 | Gm3744 (3723) | SNORD87 |
|  | **Gm3944** (3923) | SNORD111/B | Cm3808 (3787) | SNORD10 |
|  | Gm4228 (4198) | SNORD58A/B/C | Am3867 (2846) | SNORD92 |
|  | Gm4618 (4588) | SNORD91A, SNORD91B | **Cm3869** (3848) | SNORD53 |
|  | **Um4620** (4590) | SNORD72 | **Gm3944** (3923) | SNORD111/B |

**Table 2. Sensitive sites to FBL depletion.**
|  |  |  |  |
| --- | --- | --- | --- |
| **Gm4637** (4607) | SNORD121A,<br>SNORD121B | Gm4228 (4198) | SNORD58A/B/C |
|  |  | Gm4618 (4588) | SNORD91A,<br>SNORD91B |
|  |  | **Um4620** (4590) | SNORD72 |
|  |  | **Gm4637** (4607) | SNORD121A,<br>SNORD121B |

To assess whether methylation levels correlated with the expression of their corresponding SNORDs, we analyzed snoRNA abundance over the time course of FBL depletion. This was achieved by extracting deep sequencing read counts for individual snoRNA species from the RiboMethSeq dataset (**Table S7**). While most H/ACA snoRNAs remained unaffected following FBL silencing, we observed significant downregulation of 20 C/D box snoRNAs after 48 h of FBL depletion (**Figure 4I** and **Table 3**). At 72 h, only three SNORDs were significantly downregulated (**Table 3**). The lack of major changes in SNORD expression levels over time suggests that FBL depletion had no time-dependent impact on SNORD expression or stability (**Figure 4J)**. Additionally, we identified a large group of SNORDs that appeared to be resistant to FBL silencing **(Figure 4K**). GO analysis of the host genes for downregulated C/D box snoRNAs revealed categories related to ribosomal small subunit biogenesis, cytoplasmic translation, and small ribosomal subunit (SSU)-rRNA maturation from the tricistronic rRNA transcript, underscoring the significance of these host genes in translation-related mechanisms (**Figure 4L**). However, consistent with previous studies ^60,61^, we did not observe a direct correlation between sites with significantly decreased Nm levels (FC ≥ 1.3) and the intracellular abundance of their guiding SNORDs (**Table 3**), suggesting that additional factors may influence Nm site stability.

**Table 3.** SNORDs downregulated upon FBL silencing. Analysis of RiboMethSeq data for differential expression of snoRNAs revealed 20 SNORDs to be downregulated at 48 h and 23 SNORDs at 72 h upon FBL KD. In bold are highlighted the SNORDs data guide for significantly downregulated Nm sites upon FBL KD.

| Downregulated<br>snoRNAs at 48 h | Nm site | Downregulated<br>snoRNAs at 72 h | Nm site |
| --- | --- | --- | --- |
| SNORD104 | 28S-Cm1340 | SNORD117 | 18S-1320 (predicted) |
| SNORD117 | 18S-1320 (predicted) | SNORD83A | 18S-Am468 |
| SNORD41 | 28S-Um4306 | SNORD41 | 28S-Um4306 |
| SNORD69 | 28S-Gm4494 | SNORD13 | 18S-1322 (predicted) |
| SNORD13 | 18S-1322 (predicted) | SNORD3A | unknown |
| SNORD3A | unknown | SNORD69 | 28S-Gm4494 |
| SNORD6 | 28S-Gm2424 | SNORD104 | 28S-Cm1340 |
| SNORD82 | 18S-Am1678 | SNORD82 | 18S-Am1678 |
| SNORD83A | 18S-Am468 | SNORD6 | 28S-Gm2424 |
| SNORD15B | 28S-Am3785 | SNORD105B | 18S-Um799 |
| SNORD17 | 28S-Um3818 | SNORD101 | unknown |
| SNORD46 | 28S-Am3760 | SNORD15B | 28S-Am3785 |
| SNORD11 | 18S-Gm509 | SNORD11 | 18S-Gm509 |
| SNORD14E | 18S-Cm462 | SNORD46 | 28S-Am3760 |
| SNORD105B | 18S-Um799 | **SNORD92** | 28S-Am3867 |
| SNORD38A | 28S-Am1871 | SNORD17 | 28S-Um3818 |
| SNORD59A | 18S-Am1031 | **SNORD91A** | 28S-Gm4618 |
| SNORD1B | 28S-Gm4392 | SNORD66 | 18S-Cm1272 |
| **SNORD92** | **28S-Am3867** | SNORD14E | 18S-Cm462 |
| SNORD19C | 18S-Gm683 | SNORD1B | 28S-Gm4392 |
|  |  | SNORD19C | 18S-Gm683 |
|  |  | SNORD38A | 28S-Am1871 |
|  |  | **SNORD70** | 18S-Am512 |

### FBL silencing alters ribosomal RPS28 composition

To explore how global Nm hypomethylation affects translation, we applied the RiboView analysis method to Ribo-seq data, focusing on codon usage differences across experimental conditions. Our analysis revealed no significant differences in codon usage between FBL KD and control cells (**Figure S9A**), indicating that FBL silencing did not affect translation in a codon-dependent manner. Thus, the changes in translation efficiency uncovered in the Ribo-Seq data are unlikely to result from selective pausing of specific codons. Instead, we hypothesized that the observed translational deregulation in FBL KD cells is more generalized and may stem from changes in ribosomal composition, given the role of Nm in modulating RNA-protein interactions ^62^.

To investigate whether ribosomes in FBL-depleted cells exhibit altered ribosomal protein composition, we employed differential ribosomal protein incorporation prediction by analysis of rRNA fragment (dripARF) software, a recently developed tool ^26^. This method predicts differences in ribosomal protein incorporation based on their impact on the RNase degradation patterns of residual rRNA-data typically discarded in Ribo-seq analysis. Notably, when compared with the control, both FBL KDs presented distinct patterns of ribosomal protein incorporation (**Figure 5A**). Specifically, four ribosomal proteins were significantly changed in both KDs: three from the SSU (eS17, eS28, and uS5) and one from the large ribosomal subunit (eL6) (**Figure 5A**). Because only eS28/RPS28 showed a consistent reduction in both KDs, we focused on further investigating RPS28. Using the ChimeraX software ^63^, we mapped RPS28 onto the ribosome structure 8QOI to identify nearby sites with significantly reduced Nm levels (**Figure 5B**). The 8QOI ribosome structure has a 1.9 Å resolution, the highest resolution obtained for ribosome structure to date, and provides insights into the role of rRNA modifications within the human ribosome ^64^. This analysis revealed four sites near RPS28 with decreased Nm methylation, including 18S-Gm1447, which exceeded our threshold criterion of FC ≥ 1.3. Notably, 18S-Am1678 is located very close to RPS28, and simulations on the Protein Data Bank (PDB) website suggest a potential hydrogen bond interaction between RPS28 and 18S rRNA residues 1678-1679 (**Figure 5C**).

**Figure 5.**
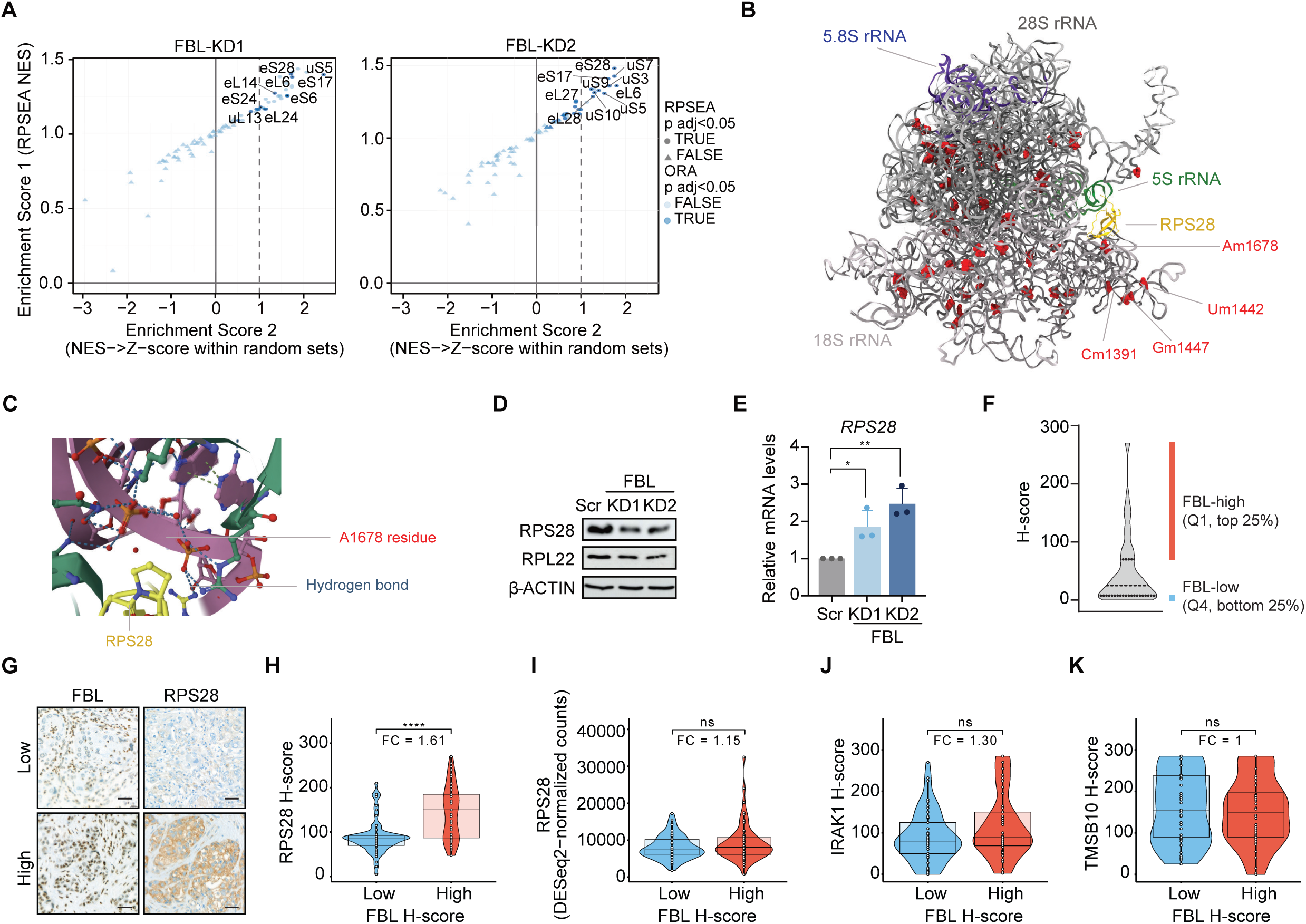
FBL depletion alters RPS28 expression and ribosomal composition, promoting the breast cancer phenotype. (**A**) Scatter plots illustrating the differential ribosomal proteins in FBL KD1 (left) and KD2 (right) compared to control. (**B)** Illustration of RPS28 (yellow) within the human ribosome, showing the significant downregulated Nm sites at 48 h of FBL depletion, generated using ChimeraX software. Nearby sites to RPS28 are labeled. (**C**) Illustration of RPS28 (yellow) forming hydrogen bond contacts with residues 1678-1679 of 18S rRNA, as captured from the PDB website. (**D**) Representative Western Blot showing the expression of RPL22 and RPS28 in control and FBL-depleted cells. β-ACTIN serves as a loading control. (**E**) RT-qPCR analysis of *RPS28* in control and FBL-depleted (KD1 and KD2) cells. The mRNA expression levels were normalized to *GAPDH*. (**F**) H-score quantification of FBL in TNBCs from SCAN-B projet. Q1 and Q4, first and fourth quartiles, respectively. (**G)** Representative IHC images showing FBL and RPS28 protein levels (low and high, respectively) in the same patient sample. Scale bar: 40 µm (**H, I**) Violin plots displaying the correlation of (**H**) FBL and RPS28 H-scores and (**I**) FBL H-score and *RPS28* mRNA in TNBC tumors from the SCAN-B project. (**J** and **K**) Violin plots displaying the correlation of FBL and (**J**) IRAK1 and (**K**) TMSB10 H-scores. Statistical analysis: All experimental data are presented as the mean ± SD, n = 3. (**E**) Unpaired two-tailed t-test, * *P* < 0.05, ** *P* < 0.01. (**H-K**) Unpaired two-tailed t-test corrected with the BH method for multiple comparisons, * adj P < 0.05, ** adj P < 0.01, *** adj P < 0.001, **** adj P < 0.0001, ns adj P > 0.05.

To further explore the interplay between FBL and RPS28, we examined the RPS28 levels following FBL silencing. FBL depletion resulted in a decrease in RPS28 protein levels, whereas RPL22, which was used as a control, remained unaffected (**Figure 5D**). Interestingly, this reduction in RPS28 protein levels occurred despite significant upregulation of *RPS28* mRNA levels (**Figure 5E**). These findings suggest the activation of compensatory mechanisms wherein cells increase RPS28 transcription to counterbalance the reduction in protein levels.

To extend our analysis, we next evaluated FBL and RPS28 protein levels using a tissue microarray (TMA) from a cohort of 227 TNBC cases within the SCAN-B study. Our findings demonstrated a significant correlation between FBL and RPS28 protein levels, though no such association was observed at the mRNA level (**Figures 5F-I)**. Similarly, high FBL levels were associated with high IRAK1 expression in the same TMA, though statistical significance was not reached, possibly due to the limited sample size (**Figures 5J).** No correlation was found between FBL and TMSB10 levels (Figure 5K). Notably, while FBL expression showed no correlation with Ki67 levels (**Figure S9B**), elevated FBL levels were linked to poorer overall survival (OS) and disease-free survival (DFS) (**Figures S9C** and **S9D**), though neither reached statistical significance due to sample size constraints and follow-up duration.

In summary, FBL depletion disrupts translation, likely by altering ribosomal composition, leading to a notable reduction in RPS28 protein. Additionally, FBL protein levels correlate with RPS28 expression in patient samples, highlighting its potential prognostic relevance.

### RPS28 promotes TNBC progression

To understand whether RPS28 plays a role in TNBC, we employed two distinct shRNAs (RPS28-KD1 and KD2) to silence its expression in MDA-MB-231 cells. Both shRNAs effectively depleted RPS28 at both the protein and mRNA levels (**Figures S10A** and **10B)**. RPS28 silencing reduced cell viability by over 50% within 72 h, impaired colony formation, and decreased wound healing capacity (**Figures 6A-D)**. Despite these effects, RPS28-depleted cells retained more than 50% of their migratory capacity (**Figures 6E-F**). Taken together, these findings suggest that RPS28 is critical for TNBC tumorigenesis.

**Figure 6.**
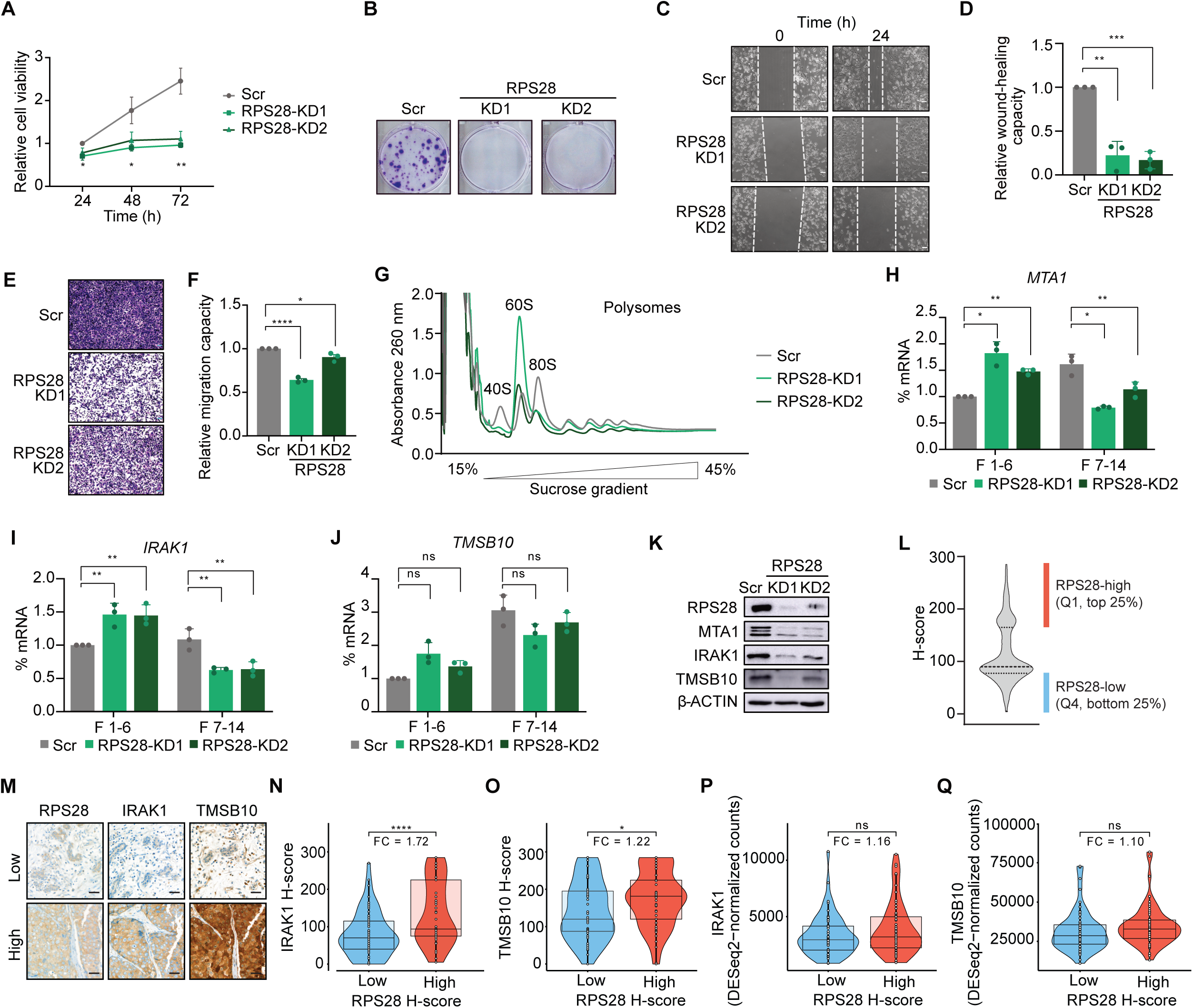
RPS28 depletion promotes breast cancer progression. (**A**) MTT assay of control and RPS28-depleted (KD1 and KD2) cells over a 3-day period (24, 48, and 72 h) upon puromycin selection. (**B**) Crystal violet staining of control and RPS28-depleted (KD1 and KD2) cells, performed 10 days after puromycin selection. (**C**) Representative images from the wound-healing assay showing the wound-healing capacity of control and FBL-depleted (KD1 and KD2) cells upon scratch generation at 0 and 24 h. (**D**) Quantification of wound-healing capacity in RPS28-depleted (KD1 and KD2) cells relative to control. Scale bar: 100 µm. (**E**) Representative images showing cells that migrated through the trans-well in Scr and RPS28-depleted (KD1 and KD2) conditions. Scale bar: 100 µm. (**F**) Quantification of the migration capacity of RPS28-depleted (KD1 and KD2) cells relative to control. (**G**) Polysome profiling for control and RPS28-depleted (KD1 and KD2) cells in a 15-45% sucrose gradient. Peaks for 40S, 60S, 80S, and polysomes are highlighted and correspond to absorbance measurements at 260 nm. (**H**-**J**) mRNA percentage distribution among the 14 fractions collected by polysome profiling for (**H**) *MTA1*, (**I**) *IRAK1*, and (**J**) *TMSB10* in control and RPS28-depleted cells. The fractions were combined as sub-polysomal (1 to 6) and polysomal (7 to 14). (**K**) Representative Western Blot showing RPS28, MTA1, IRAK1, TMSB10 protein expression in Scr control and RPS28-depleted (KD1 and KD2) MDA-MB-231 cells. β-ACTIN serves as a loading control. (**L)** H-score quantification of RPS28 in TNBCs from SCAN-B project. Q1 and Q4, first and fourth quartiles, respectively. (**M**) Representative IHC images showing RPS28, IRAK1 and TMSB10 protein levels (low and high, respectively) in the same patient sample. Scale bar: 40µm. (**N** and **O**). Violin plots displaying the correlation of RPS28 and (**N**) IRAK1 and (**O**) TMSB10 H-scores in TNBC tumors from the SCAN-B project. (**P** and **Q**) Violin plots displaying the correlation of RPS28 H-score and (**P**) *IRAK1* and (**Q**) *TMSB10* mRNAs. Statistical analysis: All experimental data are presented as the mean ± SD, n = 3. (**A, D, and F**) Unpaired two-tailed t-test, * *P* < 0.05, ** *P* < 0.01, **** *P* < 0.0001. (**H-J**) Two-way ANOVA with Dunnett’s multiple comparisons test * adj *P* < 0.05, ** adj *P* < 0.01, ns = non-significant. (**N-Q**). Unpaired two-tailed t-test corrected with the BH method for multiple comparisons, * adj *P* < 0.05, ** adj *P* < 0.01, *** adj *P* < 0.001, **** adj *P* < 0.0001, ns adj *P* > 0.05.

To determine whether RPS28 plays a role in the translation regulation of our oncoproteins of interest, we performed polysome profiling for control and RPS28 KD cells. The profiles displayed an increased peak for the free 60S ribosomal subunit and a marked reduction in peaks for the free 40S subunit and mature 80S ribosome (**Figure 6G**), which is consistent with previous studies ^65^. This suggests impairment in mature 80S ribosome formation due to defects in 40S subunit biogenesis.

RT-qPCR analysis of fractions obtained from RPS28 silenced cells showed a significantly altered mRNA distribution between subpolysomal (fractions 1–6) and polysomal (fractions 7–14) fractions for *MTA1* and *IRAK1*, with increased mRNA abundance in subpolysomal fractions and decreased abundance in polysomal fractions, suggesting impaired translation (**Figures 6H-I**). A similar trend was observed for *TMSB10*, although the changes were not statistically significant (**Figure 6J**). For *GAPDH* mRNA, no differences were observed between the control and KD cells (**Figure S10C**). At the protein level, MTA1, IRAK1, and TMSB10 decreased upon RPS28 silencing. (**Figure 6K**). A correlation with RPS28 levels and IRAK1 and TMSB10 at the protein level was also observed in TNBC from patients (**Figures 6L-O**). Notably, the mRNA levels of these transcripts remained unchanged (**Figures 6P** and **6Q**). In summary, these findings indicate that RPS28 depletion disrupts global translation by impairing mature 80S ribosome assembly and significantly altering the translation of *MTA1*, *IRAK1*, and *TMSB10* mRNAs, thereby contributing to the breast cancer phenotype.

## Discussion

Our study established FBL, the enzyme catalyzing Nm deposition, the most abundant rRNA modification, as a key driver of TNBC progression through its role in regulating protein synthesis and ribosome composition. FBL knockdown impairs tumorigenic traits, induces metabolic stress, and disrupts translation of critical oncogenes. Additionally, it reduces RPS28 protein levels and promotes ribosomal heterogeneity. The phenotypic effects of RPS28 silencing further underscored its role in FBL-mediated oncogenic translation. Analysis of human samples provides strong evidence supporting the involvement of FBL and RPS28 in this aggressive breast cancer subtype.

Consistent with previous studies, we observed significant upregulation of FBL at both the mRNA and protein levels in the TNBC cell line MDA-MB-231 compared to the normal mammary epithelial cell line hTERT-HME1 ^11,12^. Notably, recent findings suggest that while FBL is typically a marker of poor prognosis in early-stage breast cancer, a subset of aggressive tumors (approximately 10% of cases) displays reduced FBL expression, potentially disrupting ribosome biogenesis ^66^. This indicates that a low ribosomal count may be a characteristic feature of these specific tumor types. FBL depletion markedly reduced proliferation, colony formation, and migration of MDA-MB-231 cells, underscoring its critical role in the breast cancer phenotype. As an essential gene, the complete loss of FBL is lethal ^67^, and the observed reduction in proliferation is expected due to the high knockdown efficiency. Notably, calcein assays demonstrated that FBL KD cells were viable under our experimental conditions.

Several studies have suggested that modulation of Nm levels plays a key role in translational regulation ^68–70^. In this study, we showed that FBL depletion significantly reduced ribosome availability and impaired global translation. Additionally, FBL knockdown resulted in the accumulation of stalled pre-initiation complexes, specifically half-mer polysomes, indicative of defects in ribosome biogenesis ^46,47^. Given the reduction in translation efficiency and the associated metabolic disturbances, we assessed the bioenergetics of our cellular model. Notably, we observed significant suppression of glycolysis in FBL-depleted cells. Glycolysis, the primary bioenergetic pathway for energy production in cancer cells, often referred to as the Warburg effect, is frequently upregulated, despite its low energy efficiency ^71^. Targeting glycolytic metabolism is a critical mechanism for tumorigenesis and the acquisition of stem cell-like properties in breast cancer ^72,73^, and the impact of FBL on glycolysis further underscores its importance in TNBC metabolism and progression.

Furthermore, FBL silencing reduced the translation efficiency of oncogenes, such as *MTA1*, *IRAK1*, and *TMSB10*, with this effect observed in the TNBC cell line but not in normal mammary epithelial cells. MTA1 promotes key tumorigenic processes, including angiogenesis, proliferation, and epithelial-to-mesenchymal transition, in multiple cancers, including breast cancer ^74^. IRAK1 drives TNBC metastasis, recurrence, and paclitaxel resistance ^56^. Additionally, TMSB10 is linked to cell invasion and metastasis in various cancers, including TNBC ^58,75^, and its presence in the serum of patients with breast cancer makes it a potential diagnostic biomarker ^58^.

Using the quantitative deep-sequencing-based technique RiboMethSeq, we remapped all rRNA Nm sites in the MDA-MB-231 and hTERT-HME1 cell lines. Most sites (86/110, 78.2%) were identified as stable, indicating their essential role in maintaining ribosomal function. In contrast, variable sites (24/110 = 21.8%) are likely candidates for dynamic regulation during cellular transformation, contributing to ribosomal heterogeneity and potentially supporting specialized translational functions ^18^. Among these variable sites, 18S-Am576 was hypermethylated in a patient cohort of TNBC samples compared to that in healthy tissues ^22^. We also showed that Nm is more sensitive to FBL depletion at some specific sites (20 sites after 48 h of FBL depletion and 26 after 72 h). Although it has been proposed that sites hypersensitive to FBL depletion are predominantly modified sub-stoichiometrically ^60^, our findings provide limited support for this trend. Instead, site-specific rRNA methylation appears to be influenced by multiple factors, including differential accessibility of residues to snoRNPs, FBL levels, and pre-rRNA folding during ribosomal assembly ^18^.

As previously reported ^18,60^, we observed no strict correlation between the expression levels of snoRNAs targeting Nm sites and QuantScores. Similarly, there was no clear direct relationship between the sites whose Nm levels were particularly sensitive to FBL depletion and the intracellular abundance of the corresponding guiding snoRNAs. This lack of correlation does not appear to be due to technical limitations, as advanced sequencing approaches such as TGIRT-Seq in *Drosophila* have yielded similar findings ^76^.

SNORD92, SNORD91A, and SNORD70, which guide Nm modifications that were diminished and concurrently downregulated after 72 h of FBL depletion, could represent promising therapeutic targets. For instance, SNORD92 is upregulated in patients with TNBC ^77^, and SNORD91A is an exosome-associated molecule known to induce local immune microenvironment remodeling, influencing the progression of esophageal squamous cell carcinoma ^78^. However, no information is currently available on SNORD70, highlighting it as a new area for exploration. Moreover, the absence of a global correlation between snoRNA levels and their corresponding Nm rRNA, along with the relative abundance of orphan snoRNAs in humans, suggests that additional non-canonical functions contributing to breast tumorigenesis should be revealed in future studies.

Ribosomal heterogeneity also arises from changes in the ribosomal protein composition^79^. FBL depletion reduces RPS28 protein levels, a ribosomal protein that crosslinks with mRNA at the E-site of the ribosome ^80^. RPS28 forms hydrogen bonds with two rRNA nucleotides, including Nm-marked 18S-Am1678, which becomes hypomethylated upon FBL silencing. Given that a decrease in methylation at 18S-Gm1447 leads to loss of the nucleolar protein LYAR ^17^, a similar effect on RPS28 at Am1678 is plausible. RPS28 is located near the -5 and -6 mRNA regions ^81^, which are involved in translation initiation in non-AUG codons ^82^, and is implicated in IRES-mediated translation initiation ^83^, suggesting its broader role in translation regulation.

Similar to FBL silencing, RPS28 depletion reduced translation and protein levels of MTA1, IRAK1, and TMSB10, impairing the tumorigenic properties of breast cancer cells. Given its proposed role as a prognostic factor in osteosarcoma ^84^, RPS28 may play a similar role in breast cancer by regulating the translation of these oncogenes. RPS28 also has an extra-ribosomal function in pre-rRNA processing, and its silencing causes the accumulation of 45S and 30S pre-rRNAs, leading to ribosome biogenesis defects ^65,82^. However, because we did not assess the impact of RPS28 on pre-rRNA processing in our model, we cannot rule out that these defects contribute to the translational defects of MTA1, IRAK1, and TMSB10, as FBL is also involved in pre-rRNA processing.

In conclusion, our study highlighted FBL as a critical regulator of translation and ribosome composition in TNBC. FBL depletion impairs the translation of oncogenes, such as MTA1, IRAK1, and TMSB10, thus reducing tumorigenic properties. Additionally, we revealed the pivotal role of RPS28 in driving the TNBC phenotype, which was influenced by FBL. These findings offer new insights into TNBC progression of TNBC and potential therapeutic targets.

## Limitations of our study

Our study did not investigate the non-canonical role of FBL in TNBC. Beyond its methyltransferase activity, FBL drives cancer cell proliferation and increases sensitivity to DNA-damaging agents ^85^. It also contributes to histone H2AQ104 methylation, a modification specifically enriched in the 35S rDNA transcriptional unit ^86^. Examining these non-canonical functions could provide valuable insights into the broader role of FBL in TNBC.

In contrast to its well-established role in rRNA, the function of Nm in internal mRNA sites remains poorly understood. Nanopore sequencing identified Nm modifications in *MTA1*, *IRAK1*, and *TMSB10* mRNAs ^87^, which appear to be *bona fide* FBL targets based on PAR-CLIP data ^88^. Since Nm at internal mRNA sites is known to enhance mRNA stability, future studies investigating how FBL silencing affects the stability and regulation of these oncogenic mRNAs could significantly expand our findings.

## Acknowledgments

We would like to thank the Aguilo Lab members for their useful discussions. We acknowledge the Biochemical Imaging Centre at Umeå University and the National Microscopy Infrastructure, NMI (VR-RFI 2019-00217) for providing assistance with microscopy. We are thankful to Doris Lindner for the technical support. This research was supported by grants from the Knut and Alice Wallenberg Foundation, Umeå University, Västerbotten County Council, Swedish Research Council (2017-01636; 2022-01322), Cancerfonden (190337 Pj; 22 2455 Pj), and Kempe Foundation (JCK-2150). ME-S and CP are part of the ROPES ITN, which received funding from the European Union’s Horizon 2020 research and innovation program under Marie Sklodowska-Curie grant agreement number 956810. Grants from the Deutsche Forschungsgemeinschaft (DFG, German Research Foundation) project number 439669440 TRR319 RMaP TP A06 were given to the F.T. The Seahorse platform was supported by grants from the Kempe Foundation (JCK-1526) and the Västerbotten County Council (ALF) equipment grant (VLL-504771).

## Author contributions

FA conceived and supervised the study. PG, KK, ME-S, and CW performed experiments. JS, Eliana D, CP, and ED performed bioinformatics analysis. VM and YM performed the RiboMethSeq and data analysis. JDG assisted with the design and interpretation of seahorse assays. AM performed the total proteomics and analysis. PW, RW, and AB-C assisted with human sample experiments and analysis. BP analysed and quantified the IHC staining. FT assisted with the Ribo-seq experiment and data analysis. PG and FA wrote the manuscript. All authors have reviewed and edited the manuscript.

## Data availability

The mass spectrometry proteomics data were deposited in the ProteomeXchange Consortium via the PRIDE partner repository with the dataset identifier PXD051268. Ribo-seq data are available in the GEO database under GEO accession number GSE273078. RiboMethSeq data can be accessed in the ENA under ID PRJEB75039.

## Declaration of generative AI and AI-assisted technologies in the writing process

During the preparation of this study, the authors used Open AI’s ChatGPT to refine the writing process and improve the readability and language of the manuscript. After using this tool/service, the authors reviewed and edited the content as needed, and took full responsibility for the content of the published article.

## Conflict of Interest Disclosure

The authors have no conflict of interest to disclose.

## Notes

### Competing Interest Statement

The authors have declared no competing interest.

### Summary of Updates

The figures and supplemental information were missing in the former manuscript.

