## Supplementary information for "Fibrillarin shapes oncogenic protein pools and ribosomal composition in triple-negative breast cancer"

**Supplementary figures:**

**Figure S1.** Viability and apoptosis in FBL-depleted cells.

**Figure S2.** Expression of 28S and 18S rRNA upon FBL depletion and quality control for Ribo-Seq analysis.

**Figure S3.** FBL depletion leads to differential expression of transcripts related to cell growth and migration.

**Figure S4.** mRNA distribution among polysome fractions.

**Figure S5.** Comparison of Nm levels in hTERT-HME1 and MDA-MB-231 cells.

**Figure S6.** Evaluation of Nm levels in control and FBL KD1 MDA-MB-231 cells at 48 h after FBL silencing.

**Figure S7.** Evaluation of Nm levels in control and FBL KD1 MDA-MB-231 cells at 72 h after FBL silencing.

**Figure S8.** Quantification of Nm levels in FBL KD1 cells after 72 h of FBL depletion and identification of FBL sensitive sites.

**Figure S9.** Codon enrichment in FBL-KD cells and *GAPDH* mRNA distribution, and expression of candidates genes in RPS28-KD cells.

**Supplementary tables:**

**Table S1. Primers and shRNA list.** Refer to the separate Excel file, Table S1.xlsx

**Table S2.** RNA-Seq data analysis in control and FBL-depleted (KD1 and KD2) cells. Data includes $Log_2$(FC) values and the corresponding adjusted P values for differential gene expression (Input/RNA-Seq). Refer to the separate Excel file, Table S2.xlsx

**Table S3.** Ribo-seq analysis in control and FBL-depleted (KD1 and KD2) cells. Data is presented as $Log_2$(FC), with the corresponding adjusted *P* values, for differential TE (RPF/RNA-Seq) and differential RPF between FBL KD1, FBL KD2, and control. Refer to the separate Excel file, Table S3.xlsx

**Table S4.** Proteomics analysis of FBL KD1 cells at 48 and 96 h after FBL depletion. Data is presented as $Log_2$(FC), with the corresponding *P* and adjusted *P* values for each time point. Refer to the separate Excel file, Table S4.xlsx

**Table S5.** Average Raw QuantScore for cell lines. The table includes the average raw QuantScore for hTERT-HME1 and MDA-MB-231 cell lines, the mean difference between the cell lines (calculated as

MDA vs. HME1), and the associated q and p values. Methylation positions for both old and new nomenclature are provided. Sites with increased methylation in MDA-MB-231 compared to hTERT-HME1 are highlighted in red, while those with decreased methylation are highlighted in blue. Refer to the separate Excel file, Table S5.xlsx

**Table S6. Average Raw QuantScore for FBL-depleted cells.** This table presents the average raw QuantScore for each position in control and FBL-depleted (KD) cells at 48 and 72 hours. Methylation positions are provided using both old and new nomenclature. Refer to the separate Excel file, Table S6.xlsx

**Table S7. Differential snoRNA expression**. Data is presented as Log2(FC) with corresponding P and adjusted P values for 48 and 72 hours of FBL depletion. Results for each time point are provided in separate sheets. Refer to the separate Excel file, Table S7.xlsx

### Figure S1

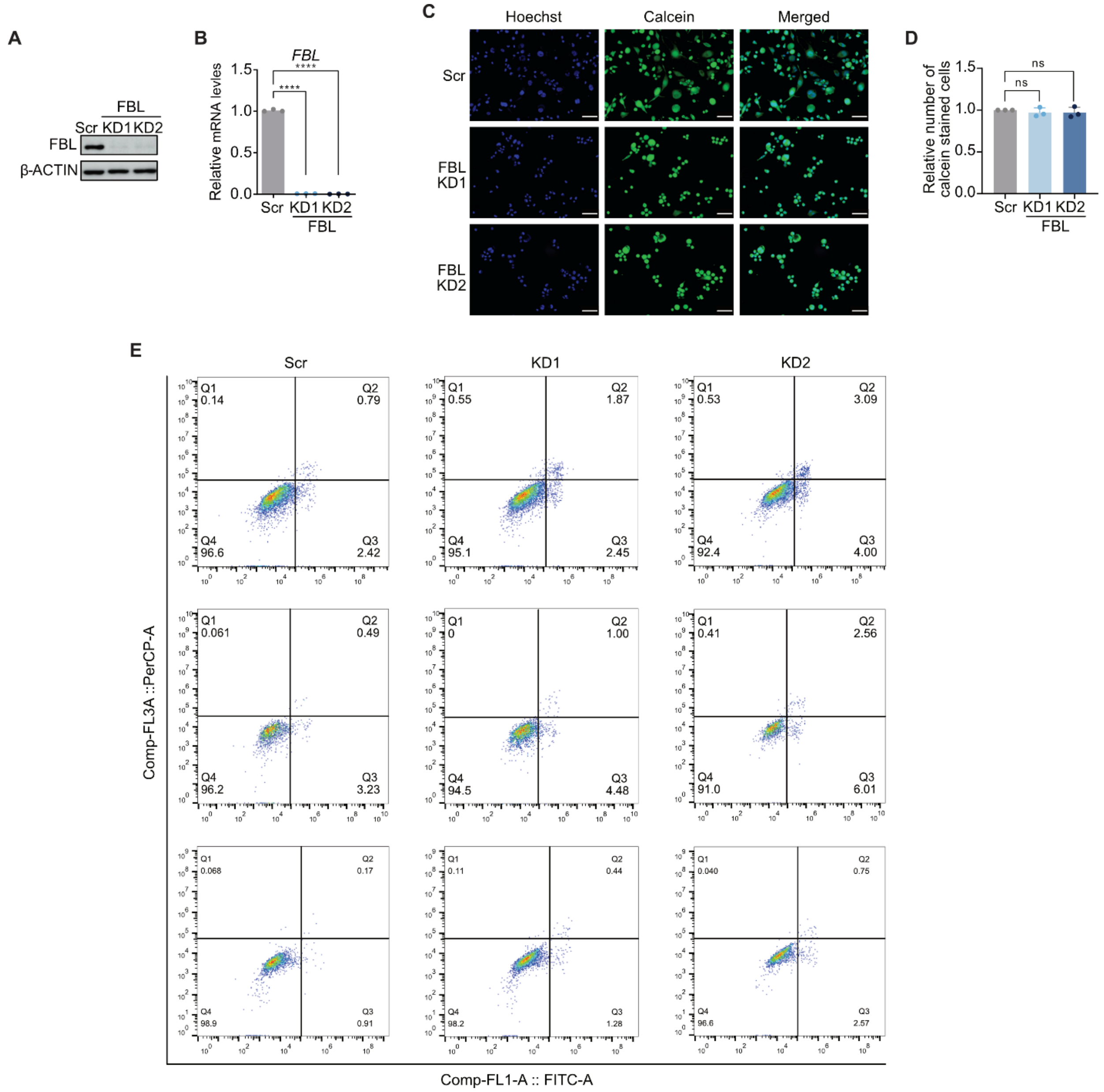


**Figure S1. Viability and apoptosis in FBL-depleted cells.** (**A**) Representative Western Blot for FBL expression in control and FBL-depleted (KD1 and KD2) MDA-MB-231 cells. β-ACTIN serves as a loading control. (**B**) RT-qPCR analysis of *FBL* in control and FBL-depleted (KD1 and KD2) cells. The mRNA expression levels were normalized to *GAPDH* (**C**) Representative images for calcein AM staining of control and FBL-depleted cells (KD1 and KD2). Calcein is marked with green, and nuclei are stained with Hoechst 33342 and depicted in blue. Scale bar: 100 µm. (**D**) Column plot depicting the relative number of calcein-stained cells in control and FBL-depleted cells (KD1 and KD2). (**E**) Flow cytometry scatter plots depicting the number of cells stained with 7ADD and Annexin V in control and FBL-depleted cells (KD1 and KD2). The Y-axis shows 7ADD (red) staining (Comp-FL3A::PERCP-A), and the X-axis shows Annexin V staining (green) (Comp-FL1-A::FITC-A).

Statistical analysis: (**B**) Paired one-tailed t-test, **** $P < 0.0001$, (**D**) Paired two-tailed t-test, ns = non-significant. Data is presented as mean ± SD, n = 3.

### Figure S2

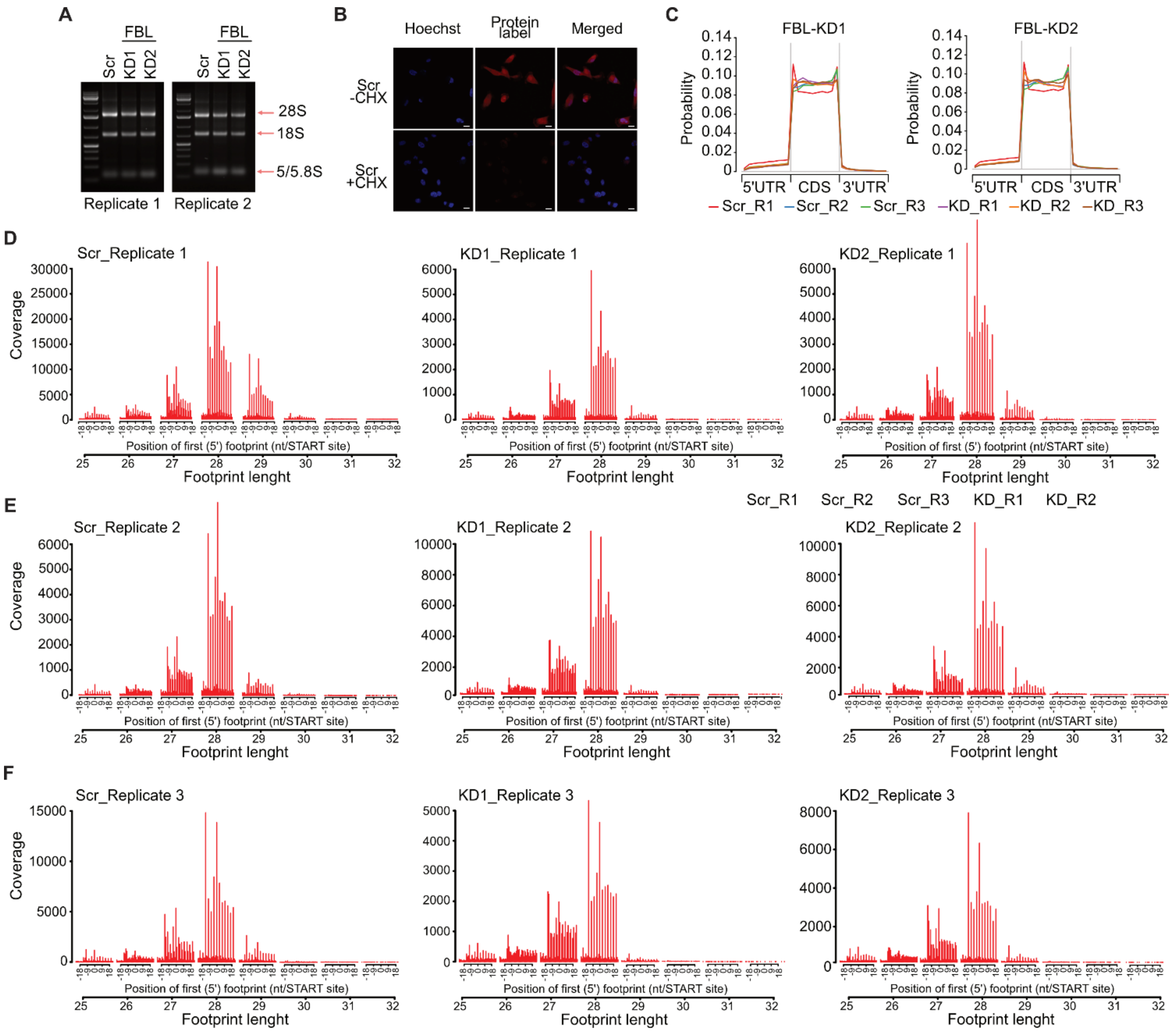


**Figure S2. Expression of 28S and 18S rRNA upon FBL depletion and quality control for Ribo-Seq data.** (**A**) Representative images of agarose gels depicting the 5/5.8S, 28S, and 18S in control and FBL-depleted (KD1 and KD2) cells from two independent biological replicates. (**B**) Representative images of nascent proteins labeled in Scr control and Scr control cells treated with CHX. Hoechst 33342 (blue) was used to mark the nuclei. The protein label solution labeled *de novo* peptides (red). Scale bar: 10 µm. (**C**) Metagene plot showing the restriction of footprints in gene coding regions for FBL-KD1 (left) vs. control and KD2 vs. control (right). (**D**-**F**) Bar plots representing the coverage obtained from different footprint lengths for all 3 biological replicates.

### Figure S3

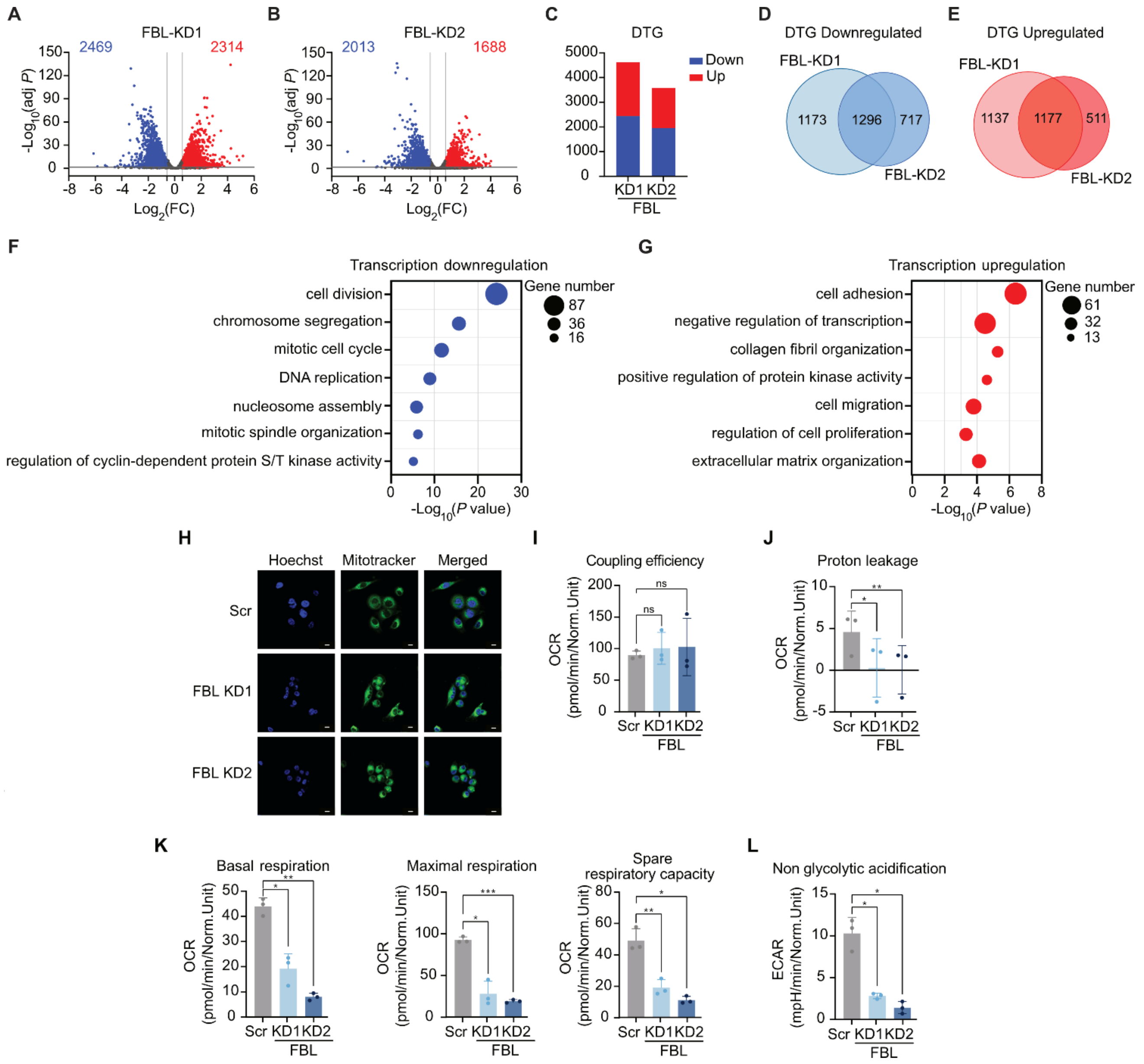


Figure legend on the next page

**Figure S3. FBL depletion leads to differential expression of transcripts related to cell growth and migration. (A** and **B)** Volcano plot illustrating differentially transcribed genes (DTG) in FBL (**A**) KD1 and (**B**) KD2 compared to Scr control. (**C**) Column plot depicting DTG genes for FBL-KD1 and KD2 that undergo downregulation (blue) or upregulation (red). (**D** and **E**) Venn diagram depicting the overlap between FBL-KD1 and KD2 for (**D**) downregulated and (**E**) upregulated DTG. (**F** and **G)** Gene ontology (GO) analysis of biological processes associated with common (**F**) downregulated and (**G**) upregulated DTG in FBL-depleted (KD1 and KD2) cells. (**H**) Representative images of mitochondria labeling with MitoTracker$^{TM}$ (green) in control and FBL-depleted cells. Hoechst 33342 (blue) was used to mark the nuclei. Scale bar: 10 µm. (**I** and **J**) Assessment of (**I**) coupling efficiency and (**J**) proton leakage in control and FBL-depleted (KD1 and KD2) cells based on the OCR. (**K**) Basal respiration, maximal respiration, and spare respiratory capacity in control and FBL-depleted (KD1 and KD2) cells. (**L)** Glycolysis parameters in FBL-depleted (KD1 and KD2) cells based on ECAR: non-glycolytic acidification.

Statistical analysis: (**I**, **J, K** and **L**) Repeated measurements one-way ANOVA using Bonferroni's multiple comparisons test. Statistical significance was considered for * adj $P < 0.05$, ** adj $P < 0.01$, *** adj $P < 0.001$, ns = non-significant. Data is represented as mean ± SD, n = 3. The cutoff applied to RNA-Seq data analysis was $Log_2(FC) \pm 0.585$, adj $P < 0.05$.

### Figure S4

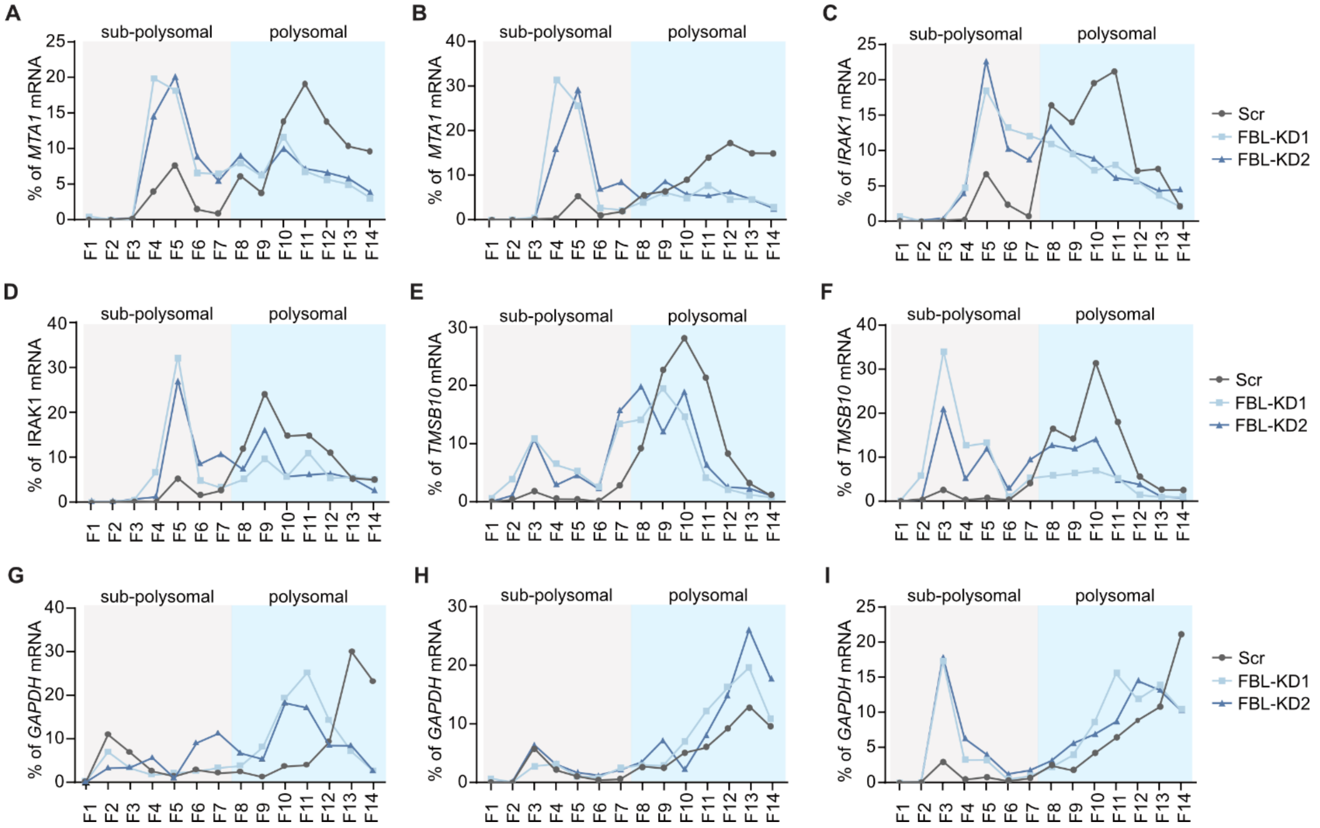


**Figure S4. mRNA distribution among polysome fractions. (A-I)** mRNA percentage distribution among the 14 fractions collected by Polysome profiling for (**A** and **B**) *MTA1*, (**C** and **D**) *IRAK1*, (**E** and **F**) *TMSB10*, and (**G-I**) *GAPDH* in control and FBL-depleted cells. Ribosomal subunits and 80S (subpolysomal) fractions are highlighted in light grey, and polysome fractions are highlighted in blue.

### Figure S5

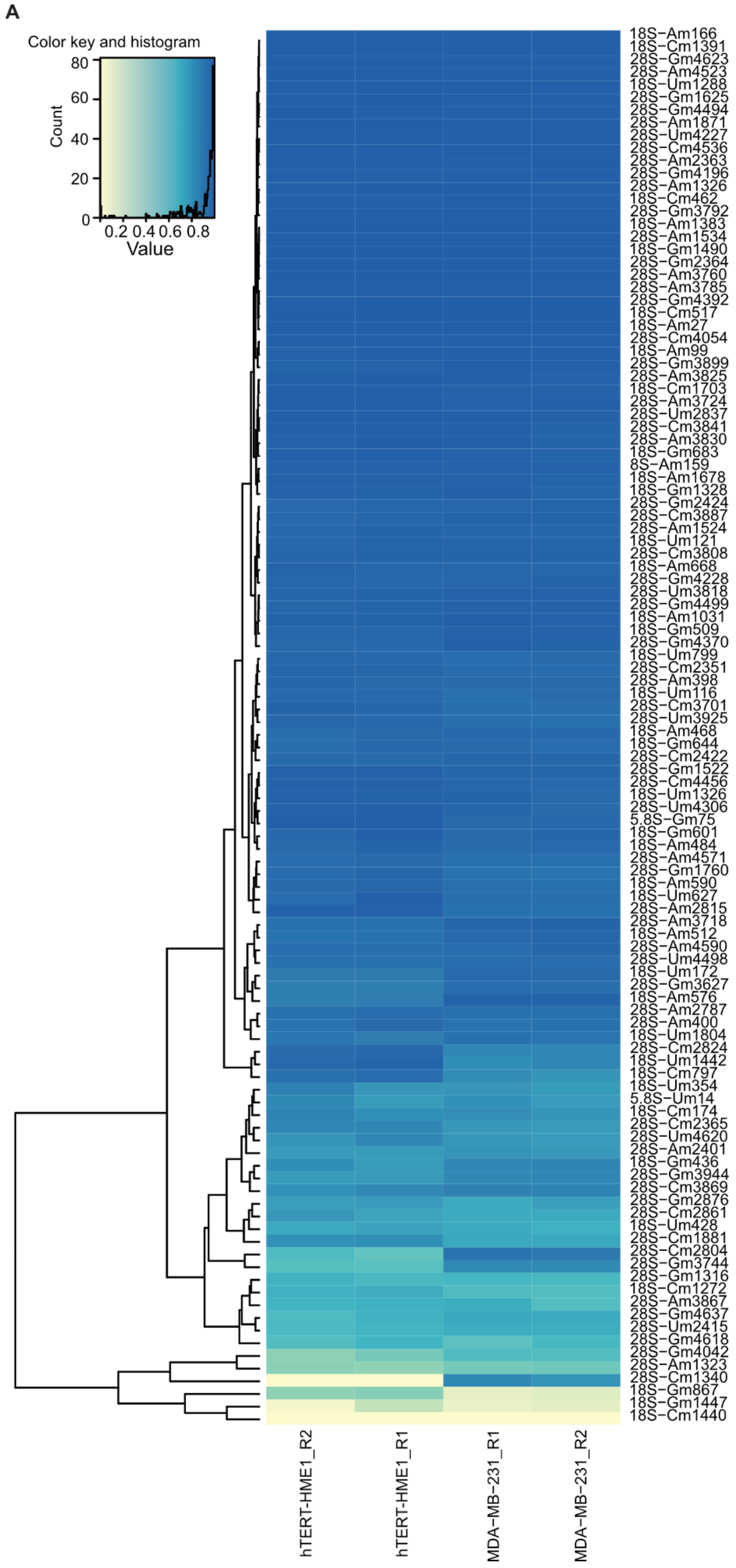


Figure legend on the next page

**Figure S5. Comparison of Nm levels in hTERT-HME1 and MDA-MB-231 cells.** (**A**) Differential QuantScore levels for 110 rRNA Nm sites were observed in two biological replicates (R1 and R2) across each cell line. The insert at the top depicts the color key, histogram, and corresponding values.

### Figure S6

A

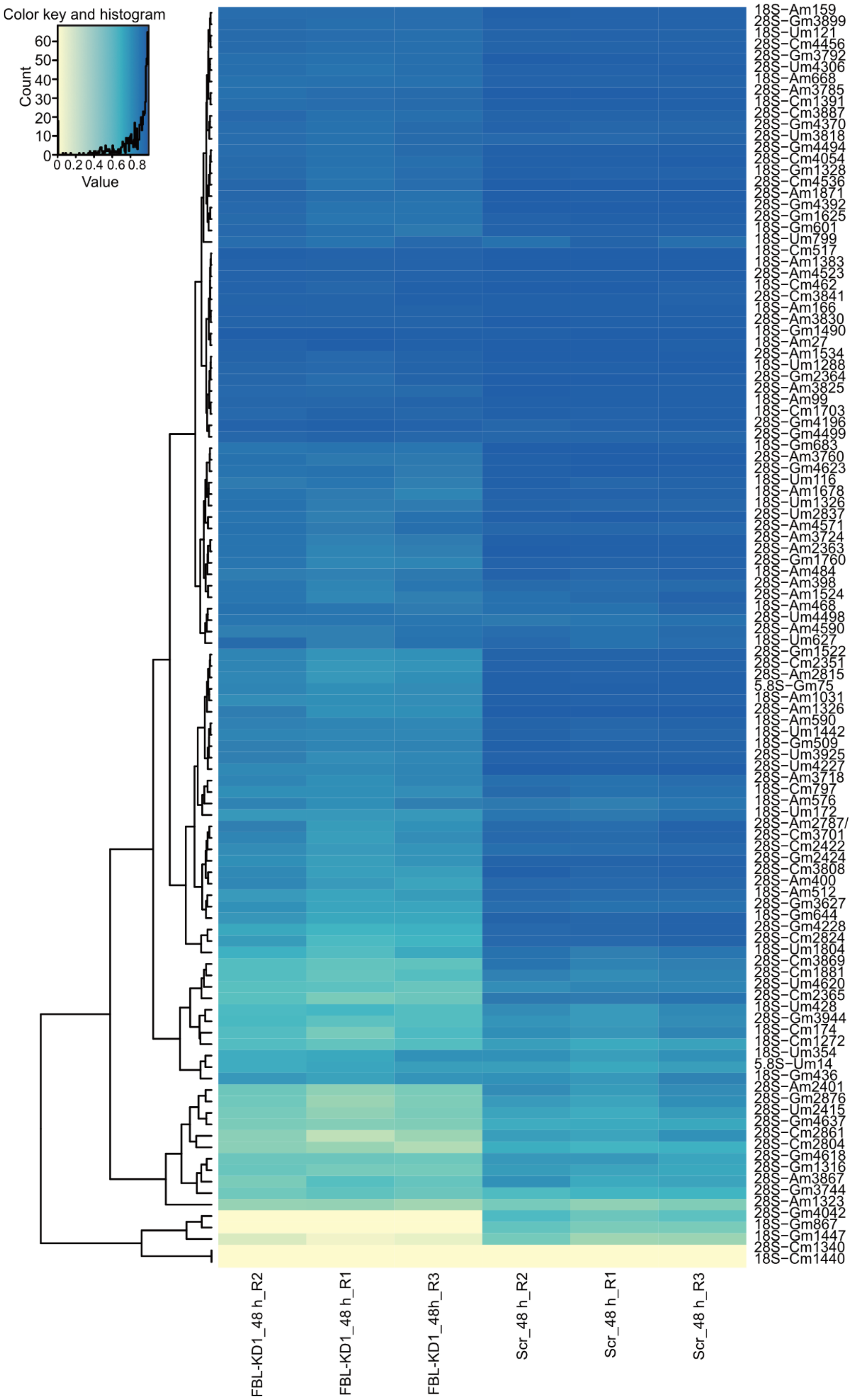


Figure legend on the next page

**Figure S6. Evaluation of Nm levels in control and FBL KD MDA-MB-231 cells at 48 h after FBL silencing.** (**A**) Differential QuantScore levels for 110 rRNA Nm sites identified in 3 biological replicates (R1-R3) across each condition. The inset at the top shows the color key, histogram, and values.

### Figure S7

A

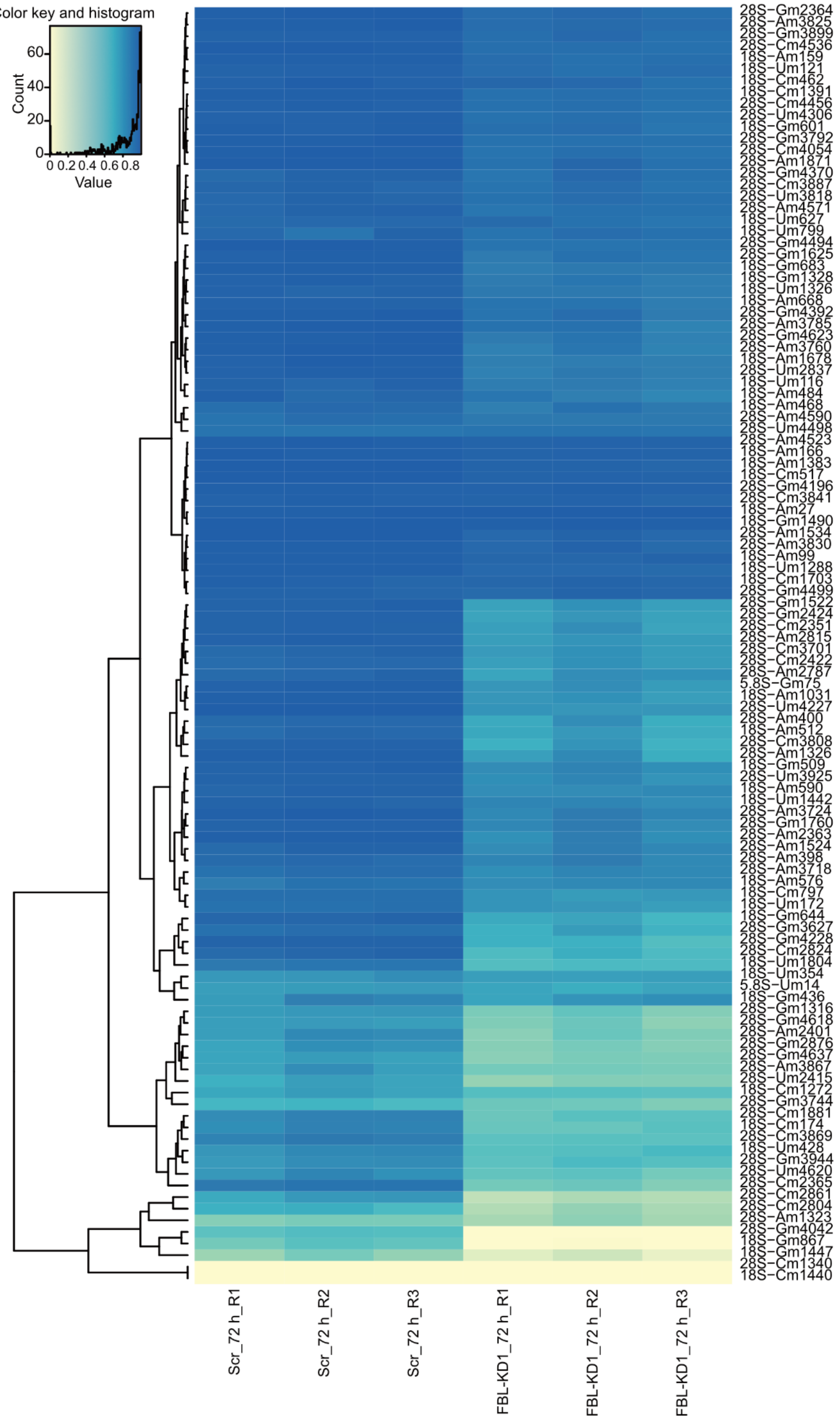


Figure legend on the next page

**Figure S7. Evaluation of Nm levels in control and FBL KD MDA-MB-231 cells at 72 h after FBL silencing.** (**A**) Differential QuantScore levels for 110 rRNA Nm sites observed in 3 biological replicates (R1-R3) across each condition. The inset at the top shows the color key, histogram, and values.

### Figure S8

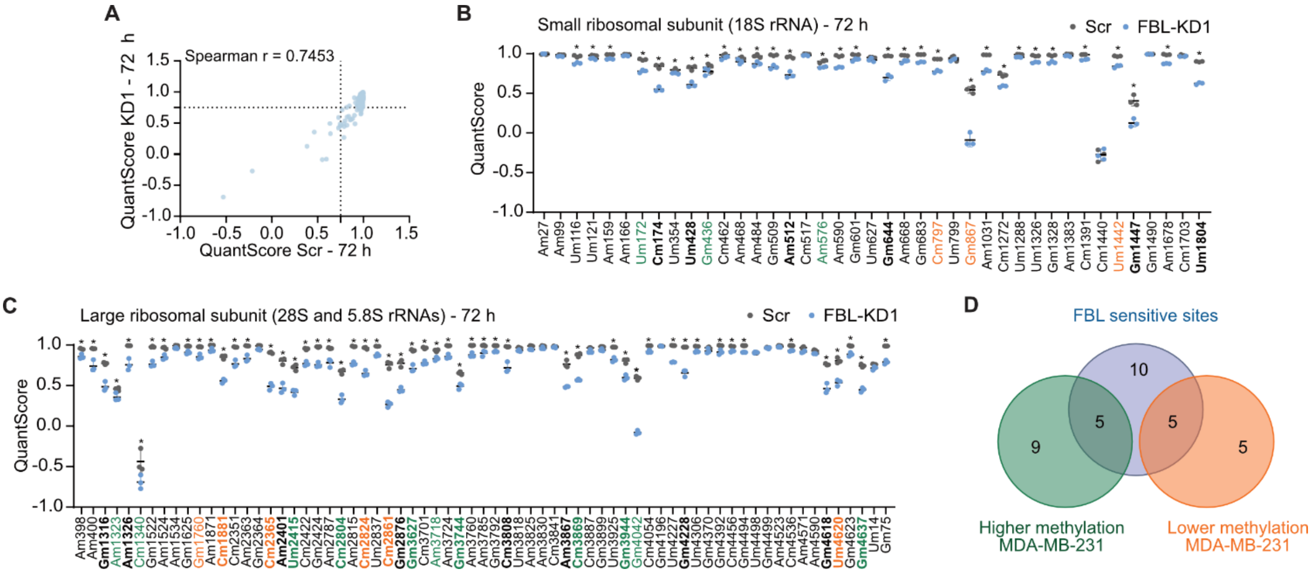


**Figure S8. Quantification of Nm levels in FBL KD1 cells after 72 h of FBL depletion.** (**A**) Correlation plot between the QuantScore from control and FBL KD1 MDA-MB-231 cells. Dot lines represent the threshold of QuantScore = 0.75, indicating fractionally methylated sites. (**B** and **C**) Dot plots illustrating the Nm levels within (**B**) the small ribosomal subunit (SSU) containing the 18S rRNA and (**C**) LSU containing the 28S and 5.8S rRNAs in control and FBL KD1 MDA-MB-231 cells at 72 h after depletion of FBL. The X-axis denotes modified nucleotide positions, and the Y-axis represents QuantScore for each site. Highlighted in green are sites with higher methylation levels, and in orange are the sites with lower methylation levels in MDA-MB-231 compared to hTERT-HME1. In bold are sites with FC ≥ 1.3 (Scr vs KD1). Data is represented as mean ± SD, n = 3. (**D**) Venn diagram depicting the overlap of differentially methylated sites in MDA-MB-231 cells compared to hTERT-HME1 cells with FBL sensitive sites displaying FC ≥ 1.3 (Scr vs KD1) at both time points after depletion of FBL (48 h and 72 h).

Statistical analysis: (**A**) Spearman correlation**,** 95% confidence interval, two-tailed *P* value, **** *P* < 0.0001. (**B** and **C**) 2way ANOVA with the two-stage linear step-up procedure of Benjamini, Krieger, and Yekutieli test for multiple comparisons. For simplicity, the significantly different sites are marked with *, regardless of the *P* value. The threshold for P value was set at less than 0.01.

### Figure S9

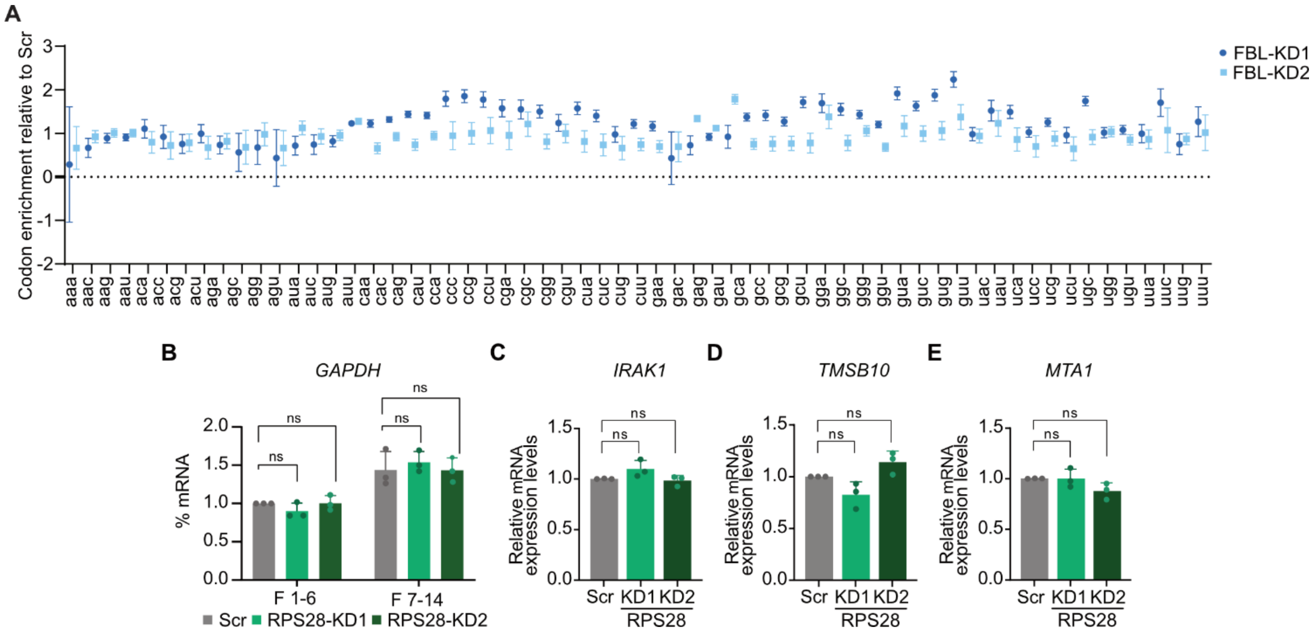


**Figure S9. Codon enrichment in FBL-KD cells and *GAPDH* mRNA distribution and expression of gene candidates in RPS28-KD cells.** (**A**) Codon enrichment analysis in FBL-depleted cells (KD1 and KD2) relative to control MDA-MB-231cells. The X-axis displays codons. The Y-axis shows the codon enrichment in FBL-KD cells relative to the control. (**B**) mRNA percentage distribution among fractions collected by polysome profiling for *GAPDH* in control and RPS28-depleted cells. The fractions were combined as sub-polysomal (1 to 6) and polysomal (7 to 14). (**C**-**E**) RT-qPCR analysis of *MTA1*, *IRAK1*, and *TMSB10* in Scr control and RPS28-depleted (KD1 and KD2) cells. The mRNA expression levels were normalized to *GAPDH*.

Statistical analysis: (**C-E**) Unpaired two-tailed t-test, ns = non-significant. Data is represented as mean ± SD, n = 3.
